# THE REGULATORY EFFECT OF LIGHT OVER FRUIT DEVELOPMENT AND RIPENING IS MEDIATED BY EPIGENETIC MECHANISMS

**DOI:** 10.1101/2020.07.30.227447

**Authors:** Ricardo Bianchetti, Nicolas Bellora, Luis A de Haro, Rafael Zuccarelli, Daniele Rosado, Luciano Freschi, Magdalena Rossi, Luisa Bermudez

**Affiliations:** Departamento de Botânica, Instituto de Biociências, Universidade de São Paulo, São Paulo, Brasil; Institute of Nuclear Technologies for Health (INTECNUS), National Scientific and Research Council (CONICET), Ruta Provincial 82, 8400 S. C. de Bariloche, Argentina; Department of Plant and Environmental Sciences, Weizmann Institute of Science, Rehovot, Israel; Instituto de Agrobiotecnología y Biología Molecular (IABIMO), CICVyA, INTA-CONICET, Argentina; Cátedra de Genética, Facultad de Agronomía, Universidad de Buenos Aires, Buenos Aires, Argentina

**Keywords:** carotenoid, chlorophyll, DNA methylation, epigenetics, fleshy fruit, phytochrome, RNA-seq, RdDM, tomato

## Abstract

Phytochrome-mediated light and temperature perception has been shown to be a major regulator of fruit development. Furthermore, chromatin remodelling via DNA demethylation has been described as a crucial mechanism behind the fruit ripening process; however, the molecular basis underlying the triggering of this epigenetic modification remains largely unknown. Here, through integrative analyses of the methylome, siRNAome and transcriptome of tomato fruits from *phyA* and *phyB1B2* null mutants, we report that PHYB1 and PHYB2 control genome-wide DNA methylation during fruit development from green towards ripe stages. The experimental evidence indicates that PHYB1B2 signal transduction is mediated by a gene expression network involving chromatin organization factors (DNA methylases/demethylases, histone-modifying enzymes and remodelling factors) and transcriptional regulators leading in the altered mRNA profile of photosynthetic and ripening-associated genes. This new level of understanding provides insights into the orchestration of epigenetic mechanisms in response to environmental cues affecting agronomical traits.

## INTRODUCTION

As sessile organisms, plants must constantly monitor their environment and continuously tune their gene expression to enable adaptation and survival (Kaiserli *et al*., 2018). Light is one of the main environmental cues that controls plant growth and development from seed germination to senescence (Galvão & Fankhauser, 2015). Plants employ different photoreceptors to detect and respond to changes in the incident spectral composition (from UV-B to far-red wavelengths), light direction and photoperiod. These photoreceptor families include (i) phytochromes (PHYs), which perceive red/far-red (R/FR) light; (ii) cryptochromes (CRYs), phototropins, and ‘Zeitlupes’, which sense blue/UV-A light; and (iii) the UV-B receptor UVR8 (Paik & Huq, 2019).

After photoreceptor activation, complex signal transduction pathways control the expression of light-regulated genes via transcriptional, posttranscriptional, and posttranslational mechanisms (Galvão & Fankhauser, 2015). Several hub proteins in the light signal transduction pathway triggered by PHYs, CRYs and UVR8 have been identified, including transcription factors (TFs) such as PHY-INTERACTING FACTORS (PIFs) and ELONGATED HYPOCOTYL5 (HY5), HY5-HOMOLOGUE (HYH), as well as the ubiquitin E3 ligase complexes comprising CONSTITUTIVE PHOTOMORPHOGENIC1 (COP1) (Galvão & Fankhauser, 2015). Both PHYA and PHYB can directly bind to target promoters (Chen *et al*., 2014; Jung *et al*., 2016) and, recently, the effect of light on alternative splicing (AS) has also been reported (Cheng and Tu 2018; Shikata et al. 2014). Furthermore, light controls protein localization through PHY-mediated alternative promoter selection, allowing plants to metabolically respond to changing light conditions (Ushijima *et al*., 2017). Finally, it is widely known that activated PHYs induce post-translational changes in PIF proteins, including sequestration, phosphorylation, polyubiquitylation, and subsequent degradation through the 26S proteasome-mediated pathway (Paik *and* Huq, 2019). Although the effect of light on plant phenotypes and the plant transcriptome has been studied for decades (Mazzella *et al*., 2005; Ibarra *et al*., 2013; Carlson *et al*., 2019), the involvement of epigenetic regulatory mechanisms in light-dependent changes in the transcriptional landscape remains poorly addressed.

Posttranslational histone modifications, such as acetylation and methylation, have been associated with the induction and repression of light-responsive genes (Perrella and Kaiserli 2016; Tessadori et al. 2009). Light-dependent enrichment of the acetylation pattern of H3 and H4 in the enhancer and promoter regions of the pea plastocyanin locus *PetE* has been reported (Chua et al. 2001), and the hyperacetylation of the *PetE* promoter is linked to the transcriptional activity of this gene (Chua et al. 2003). Moreover, a reduction in H3 acetylation is associated with a decrease in the expression of the *A. thaliana* light-responsive genes CHLOROPHYLL a/b-BINDING PROTEIN GENE (CAB2) and the RIBULOSE BISPHOSPHATE CARBOXYLASE/OXYGENASE small subunit (RBCS)(Bertrand et al. 2005). Histone methylation regulates PHY-mediated seed germination in *A. thaliana*. Upon R light illumination, photoactivated PHYB (Pfr) targets PIF1 for proteasome-mediated degradation, releasing the expression of the JUMONJI HISTONE DEMETHYLASES JMJ20 and JMJ22. As a result, JMJ20 and JMJ22 reduce the levels of H4R3me2, which leads to the activation of the gibberellic acid biosynthesis pathway to promote seed germination (Cho et al. 2012). Recently, it has been demonatrated that, in the presence of light, PHY-downstream effector HY5 recruits HISTONE DEACETYLASE 9 (HDA9) to autophagy-related genes to repress their expression by deacetylation of H3. In the darkness, HY5 is degraded via 26S proteasome and the concomitant disassociation of HDA9 leads to activated autophagy (Yang et al. 2020). Evenmore, ChIP-seq studies have revealed that many genes targeted by HY5 are enriched for specific histone marks (Charron et al., 2009).

Together with histone modification, DNA methylation is a common epigenetic mark with a direct impact on gene expression. Nevertheless, only a few reports have specifically addressed the effect of light stimuli on DNA methylation. Light-dependent nuclear organization dynamics during deetiolation are associated with a reduction in methylated DNA (Bourbousse et al. 2020). In *Populus nigra*, 137 genes were shown to be regulated by methylation during the day/night cycle (Ding et al. 2018). Moreover, photoperiod-sensitive male sterility is regulated by RNA-directed DNA methylation (RdDM) in rice (Ding et al. 2012). In tomato, plants overexpressing UV-DAMAGED DNA BINDING PROTEIN 1 (DDB1), a component of the ubiquitin E3 ligase complex, showed reduced size in reproductive organs (flowers, seeds and fruits) associated with the promoter hypomethylation and the upregulation of the cell division negative regulator *SlWEE1* (Liu et al. 2012). Finally, using a methylation-sensitive amplified polymorphism assay, DNA methylation remodelling was shown to be an active epigenetic response to different light qualities in tomato seedlings (Omidvar and Fellner 2015).

Previous studies have shown that PHYA, PHYB1 and PHYB2 are major regulators of *Solanum lycopersicum* fruit ripening and nutraceutical compounds accumulation (Gupta *et al*., 2014; Llorente *et al*., 2016; Alves *et al*., 2020; Gramegna *et al*., 2019; Bianchetti *et al*., 2018; Bianchetti *et al*., 2020). Moreover, it has also been shown that tomato fruit ripening involves epigenome reprogramming (Zhong *et al*., 2013). Here, genome-wide transcriptome, siRNAome and methylome were comprehensively analysed in fruits from *phyA* and *phyB1B2* null mutants. The results revealed that PHY-mediated gene expression modulation along fruit development and ripening involves DNA methylation regulatory mechanism.

## RESULTS

### Impact of light perception impairment on the fruit transcriptome

To investigate the role played by either PHYA or PHYB1 and PHYB2 (hereafter PHYB1B2) in overall gene expression during fruit development, the transcriptome of fruits at the immature green (IG) and breaker (BK) stages from *phyA* and *phyB1B2* null mutants as well as their wild-type (WT) counterpart, was determined by RNAseq. Among the approximately 20,000 transcriptionally active loci in each biological replicate (Supplemental Table S1), 1.2% and 2.4% at the IG stage and 9.1% and 11.2% at the BK stage were identified as differentially expressed genes (DEGs) in *phyA* or *phyB1B2* mutants, respectively, compared to the WT (Fig. 1A; Supplemental Table S2. For both genotypes, the number of exclusive DEGs was significantly lower in the IG stage than in the BK stage; similarly, the number of genes that were commonly regulated by PHYA and PHYB1B2 was 172 at the IG stage and 785 at the BK stage (Fig. 1B). Subsequently, the altered expression of approximately 76% (23/30) of the tested genes was validated by RT-qPCR (Supplemental Table S3). Comparison with previously reported expression data for genes involved in ripening regulation, ethylene biosynthesis and signalling, and carotenogenesis further validated our RNAseq data, as 90% of the analysed genes on average showed the expected mRNA profile at IG and BK stages. It is worth mentioning that most of the genes displayed the same transcript fluctuation in the WT, *phyA* and *phyB1B2* genotypes, though this was somewhat attenuated in the mutants (Supplemental Table S4). These results showed that PHY-meditated light perception regulates more genes in BK than in the early stages of fruit development and that PHYB1B2 has a more substantial impact than PHYA in the fruit transcriptome in both analysed stages. A closer look at DEGs function revealed a similar distribution of loci across MapMan categories in response to *phyB1B2* and *phyA* mutations in both developmental stages, although with distinct abundance levels (Fig. 1C). At the IG stage, eight categories were mainly represented, including at least 2% of the DEGs identified in *phyA* and *phyB1B2*: photosynthesis, lipid metabolism, phytohormone action, RNA biosynthesis, protein modification, protein homeostasis, cell wall organization, and solute transport (Fig. 1C; Supplemental Tables S5 and S6). It is worth highlighting the abundance of the DEGs within the photosynthesis category in the *phyB1B2* mutant, among which 34 out of the 37 genes were downregulated (Supplemental Table S6). In the BK stage, at least 2% of the DEGs were related to the lipid metabolism, phytohormone action, RNA biosynthesis, protein modification and homeostasis, cell wall organization and solute transport categories in both genotypes (Fig. 1C; Supplemental Tables S7 and S8). However, while *phyA* deficiency also affected carbohydrate metabolism and external stimuli (Supplemental Table S7), the *phyB1B2* mutant showed a large number of DEGs related to the cell cycle and chromatin organization (Supplemental Table S8). Interestingly, the chromatin organization category displayed 52 DEGs, 45 of which were upregulated. These genes encode nucleosome constituent histones (H3, H4, H2A and H2B); DNA methylases/demethylases; histone post-translational modifiers such as deacetylases, methylases/demethylases, histone ubiquitination factors and histone chaperones; chromatin remodelling factors; and genes involved in RNA-independent and RNA-directed DNA methylation (Supplemental Table S8). These results led us to further investigate the impact of DNA methylation on PHY-mediated gene expression reprogramming.

**Fig. 1.**
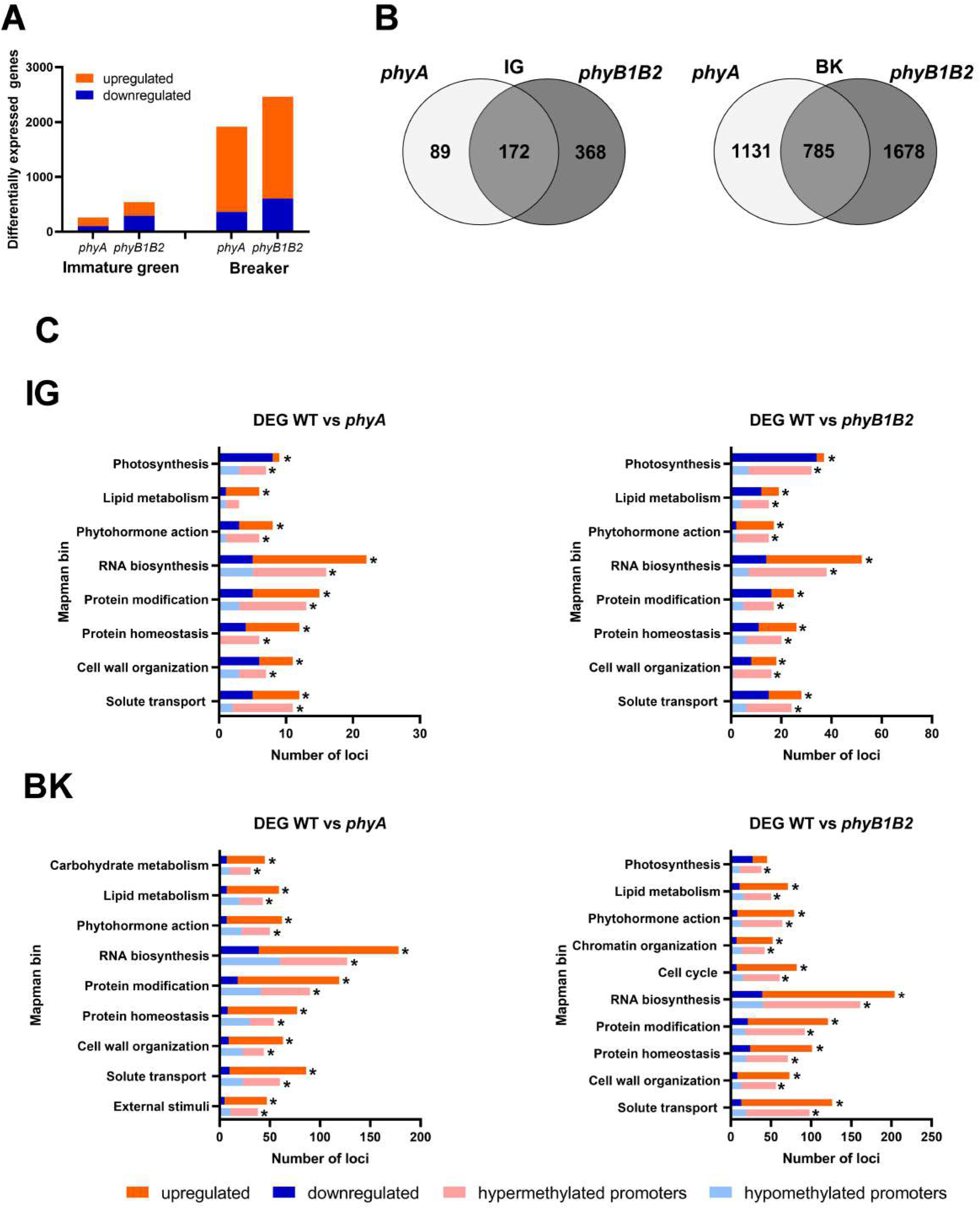
PHYA and PHYB1B2 modify the global transcriptomic profile of tomato fruit. (A) Number of differentially expressed genes (DEGs) in *phyA* and *phyB1B2* mutant fruits at immature green (IG) and breaker (BK) stages. (B) Venn diagram showing exclusive and common DEGs in *phyA* and *phyB1B2* mutants in both developmental stages. (C) Functional categorization of all DEGs and those DEGs with differentially methylated promoters (DMPs) in both analysed genotypes and stages. Only categories corresponding to at least 2% of the DEGs or DMPs in each comparison are shown (asterisks). Up-and downregulated genes are indicated in red and blue, respectively. Loci with hyper- and hypomethylated promoters are indicated in light red and light blue, respectively. DEGs and DMPs show statistically significant differences (FDR < 0.05) relative to WT.

### PHYs-dependent reprogramming of tomato fruit methylome

The global profile of methylated cytosines (mCs) in the epigenome of tomato fruits was assessed by whole-genome bisulfite sequencing in the IG and BK stages for *phyA*, *phyB1B2* and WT genotypes. In agreement with previous reports (Zhong et al. 2013; Zuo et al. 2020), regardless of the genotype and fruit stage, the greatest total number of mCs was located in the CHH context, followed by the CG and CHG contexts, while the methylation level was highest in the CG (80%) context followed by the CHG (67%) and CHH (23%) contexts (Supplemental Table S9, Supplemental Fig. S1). For further comparisons, we selected only cytosines with coverage >10X, and except for chromosome 9 in the transposable elements TEs enriched region, all samples met this cutoff. In all contexts, the highest cytosine density was associated with gene-rich euchromatic regions located at chromosome arm ends (Supplemental Fig. S1). Conversely, in symmetrical contexts (CG and CHG), the highest methylation rates were found across pericentromeric regions enriched in TEs. Yet, the highest methylation rates in CHH context was observed in gene-rich regions associated with a higher density of sRNAs (Supplemental Fig. S1), as previously reported (Corem et al. 2018). The comparison of the methylation status between the two fruit stages showed that ripening-associated demethylation (Zhong et al. 2013) occurs mainly in the CG context, especially in gene-rich regions, and that it is impaired in *phyB1B2* mutant BK fruits (Supplemental Fig. S1).

The subsequent comparison between genotypes revealed global epigenome alteration in *phy* mutants in all contexts analysed. The most remarkable observation was the presence of considerable hypermethylation in all contexts across gene-rich regions in BK-stage fruits from *phyB1B2* (Fig. 2A). In contrast, *phyA* exhibited hypermethylation in CHG and CHH contexts associated with TE-rich regions (Fig. 2A), suggesting that different PHYs control DNA methylation across specific genomic regions through distinct regulatory mechanisms. Interestingly, PHY-associated hypomethylation was exclusively detected in the CG context of gene-rich regions in IG-stage fruits from *phyA* and in the CHH context of TE-rich regions for BK-stage fruits from *phyB1B2*. In summary, these data revealed that both PHYA and PHYB1B2 affect the global methylome, but PHYB1B2 has a greater impact on ripening-associated methylation reprogramming across gene-rich genomic regions in tomato fruits. To investigate the relationship between PHY-dependent modifications in cytosine methylation and gene expression, we first identified genes with differentially methylated promoters (DMPs, 2 kb upstream the TSS) in all three contexts. Interestingly, associated with the massive alteration previously observed, the pattern of DMPs varied with the mC context, stage and genotype (Fig. 2B, Supplemental Table S10 and S11). Regarding the CG context, whereas the *phyA* mutant showed virtually the same frequency of hyper- and hypomethylated promoters in the two stages, the status of hypermethylated promoters in *phyB1B2* increased over 60% from the IG to BK stage, while the number of loci with hypomethylation decreased 50% (Fig. 2B, Supplemental Table S12). In contrast, *phyA* showed a greater number of hypermethylated promoters in the CHG context in the IG stage than in the BK stage, while the levels in the WT and *phyB1B2* mutant remained similar upon ripening (Fig. 2B, Supplemental Table S13). In the CHH context, the number of hypermethylated promoters decreased in both genotypes from the IG to BK stages (Fig. 2B, Supplemental Table S14). These results indicate that PHY deficiency results in massive promoter hypermethylation in both the IG and BK stages of tomato fruit development. Moreover, they reinforce the role of PHYB1B2 in ripening-associated demethylation and its putative effect on gene expression.

**Fig. 2.**
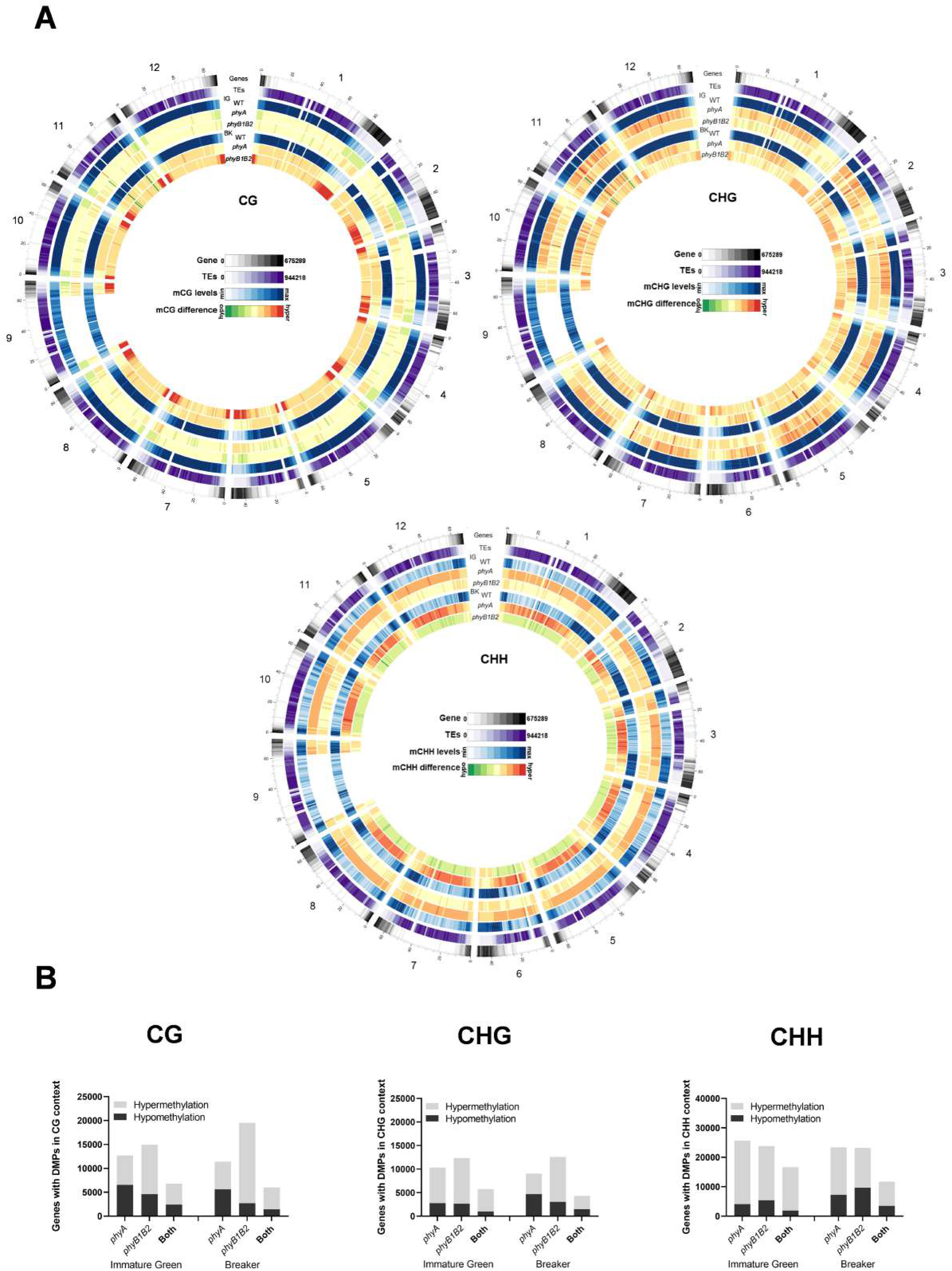
Disturbed PHYA-and PHYB1B2-dependent signalling differentially alters tomato fruit methylome. (A) Density plot of genes, transposable elements (TEs) and mC in all contexts (mCG, mCHG, mCHH) for the WT genotype. Global methylation changes for *phyA* and *phyB1B2* in comparison with the wild type (WT) at the immature green (IG) and breaker (BK) stages are shown (bin size, 1 Mb). Gene and TE densities were estimated according to the number of nucleotides covered per million. The methylation levels in the CG, CHG and CHH contexts are 40-90%, 25-80% and 10-30%, respectively. The mC difference was relative to the corresponding WT fruit stage within a −5% (hypomethylated) ≤ range ≤ +5% (hypermethylated). (B) Number of genes with differentially methylated promoters (DMPs, 2 kb upstream transcription start site) in *phyA*, *phyB1B2* and both mutants. Hyper- and hypomethylation are indicated by grey and darker-coloured bars, respectively. DMPs show statistically significant differences (FDR < 0.05) relative to WT.

### Effect of PHY-mediated differential methylation on the transcriptome

To assess whether the differential methylation of gene promoters affects mRNA levels, we crossed data from DEGs and DMPs between genotypes. Supplemental Fig. S2 shows scatter plots of promoter methylation vs mRNA fold changes for comparisons of the two genotypes at the two examined developmental stages in the three mC contexts. The most evident result was that among the thousands of loci with identified DMPs (Fig. 2B), only hundreds of the loci were also differentially expressed (Supplemental Table S15) (0.7% for IG *phyA*, 1.6% for IG *phyB1B2*, 5.6% for BK *phyA* and 7.4% for BK *phyB1B2*), raising an intriguing question about the biological significance of the extensive change in the methylation pattern observed in the mutants. In contrast, the percentages of the DEGs showing DMPs were 73% for IG *phyA*, 76% for IG *phyB1B2*, 72% for BK *phyA* and 75% for BK *phyB1B2* (Supplemental Fig. S2). Many more DEGs with DMPs were observed in BK than in IG fruits and in *phyB1B2* than in the *phyA* genotype. The functional categorization of these genes revealed a similar category distribution to the DEGs (Fig. 1C, Supplemental Tables S16-S19). At the IG stage, there were seven categories in which at least 2% of the loci showed DMPs and differential expression in both genotypes: photosynthesis, phytohormone action, RNA biosynthesis, protein modification and homeostasis, cell wall organization and solute transport, whereas *phyB1B2* additionally impacted lipid metabolism (Fig. 1C). In the BK stage, the categories in which at least 2% of the DEGs showed DMPs were lipid metabolism, phytohormone action, RNA biosynthesis, protein modification and homeostasis, cell wall organization and solute transport-related functions in both genotypes, while only *phyA* impacted carbohydrate metabolism and external stimuli, and only *phyB1B2* affected photosynthesis, chromatin organization and cell cycle categories.

Interestingly, when comparing IG and BK stages, 42.5%, 34.2% and 18.8% of the DMPs were associated with DEGs, while 79.5%, 76.6% and 71.5% of the DEGs showed differences in promoter methylation in WT, *phyA* and *phyB1B2*, respectively (Supplemental Fig. S3). These results demonstrate that the altered mRNA profile of *phyA* and *phyB1B2* fruits are associated with marked changes in promoter methylation; however, massive genome-wide PHY-induced methylation reprogramming has a still uncharacterized role beyond the regulation of mRNA accumulation. Moreover, promoter methylation has a greater effect on gene expression regulation during BK than in the IG stage. Additionally, the data showed that PHYB1B2 has a more extensive influence on gene expression regulated via promoter methylation than PHYA, reinforcing the above conclusion that PHYB1B2 affects CG ripening-associated demethylation (Supplemental Fig. S2).

### The sRNAome is altered by PHY deficiency

To assess the involvement of RdDM in PHY-mediated transcriptome regulation, the sRNAome was analysed in fruits at the IG and BK stages from both mutants and the WT genotype (Supplemental Table S20A). A total of 28,314 clusters of sRNAs were identified across the whole genome in at least one of the samples, including 7,984 in gene bodies, 7,863 in promoter regions, 7,966 in TEs and the remaining 4,501 across intergenic regions (Supplemental Fig. S1, Supplemental Table S20B). The methylation level was evaluated for each sRNA cluster-targeted genomic region (sCTGR) and, as previously observed for promoter regions, a higher proportion of hypermethylation was observed in BK fruits from *phyB1B2* in the CG symmetrical context. Moreover, the greatest number of differentially methylated sCTGRs was observed in the asymmetrical context CHH (Fig. 3A, Supplemental Table S20G-J).

**Fig. 3.**
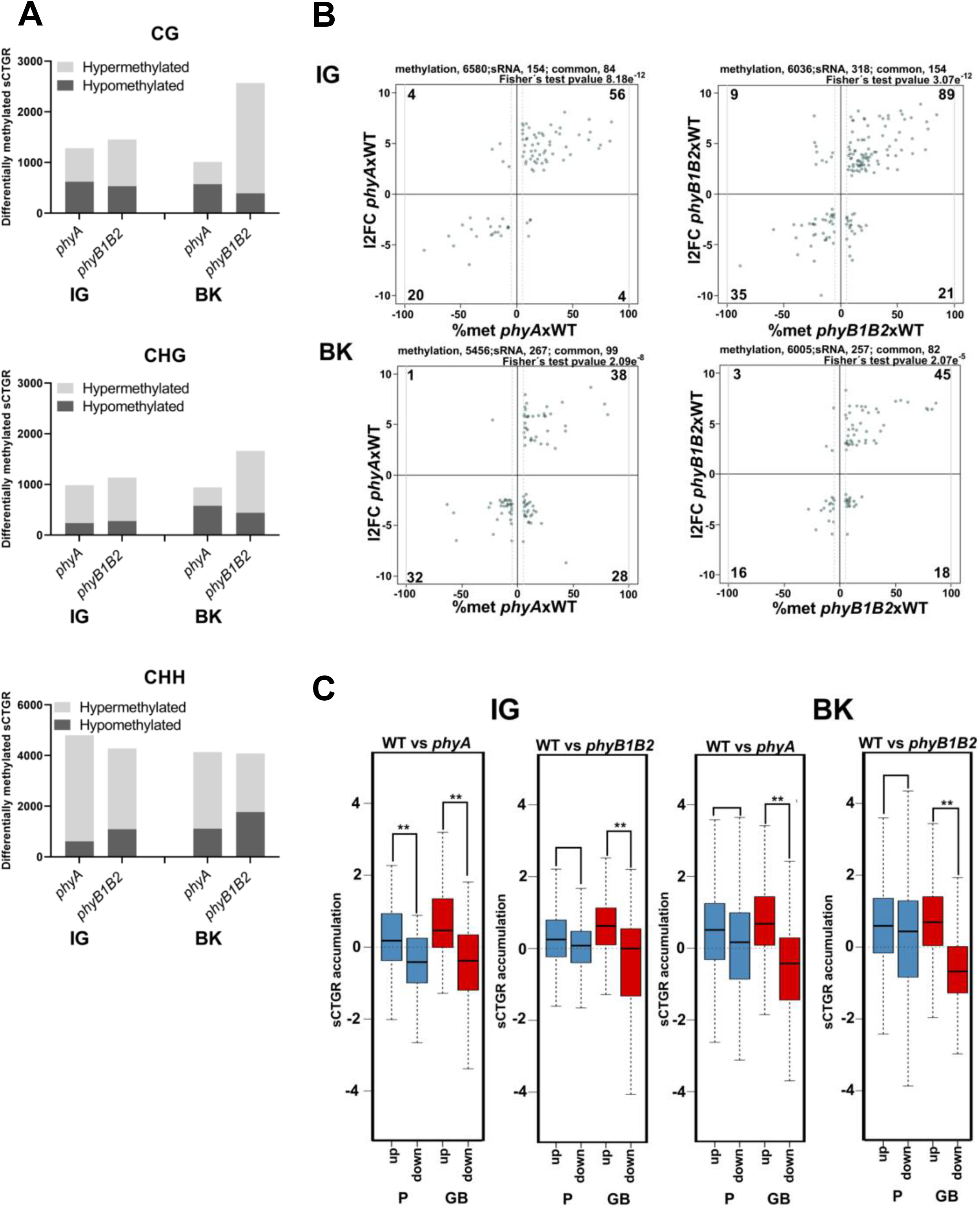
Phytochrome deficiency impacts the sRNAome profile. (A) Total number of differentially methylated sRNA cluster-targeted genome regions (sCTGRs). (B) Scatter plots show the relationship between the differential accumulation of cluster sRNAs and a minimum of 5% differential methylation of their sCTGRs. The result of Fischer′s test for the association of the two datasets is shown (*p*≤ 2.07e^−5^). (C) Boxplots show changes in the accumulation of cluster sRNAs in promoter (P, 2 Kb upstream of the 5’ UTR end) and gene body (GB) regions for up- and downregulated DEGs. Asterisks indicate statistically significant differences by the Wilcoxon–Mann–Whitney test (** *p*<0.0001). All results represent the comparison of *phyA* and *phyB1B2* to the wild type in immature green (IG) and breaker (BK) fruit stages.

sCTGR methylation levels and sRNA accumulation data were intersected, and among a total of 154, 318, 267 and 257 differentially accumulated sRNA clusters (Supplemental Table S20C-F), 84, 154, 99 and 82 also showed differential methylation in their targeted genomic region for *phyA* IG, *phyB1B2* IG, *phyA* BK and *phyB1B2* BK fruits, respectively (Fig. 3B, Supplemental Table S20G-J), showing a strong association (*P*<0.005) between the two datasets. Intriguingly, this positive association was not observed in the transition from the IG to BK stages (Supplemental Fig. S4), suggesting that the global methylation changes via RdDM could be attributed to PHY deficiency. Moreover, a clear disturbance in sRNA accumulation was observed in *phyB1B2*, since almost no clusters with less sRNA accumulation were observed in BK compared to the IG stage (Supplemental Fig. S4).

Further, we analysed whether this association between sRNA accumulation and sCTGR methylation impacted gene expression levels. Notably, regardless of the fruit developmental stage, changes in the accumulation of sRNA located in gene bodies (GBs), and not in the promoter (P) region, were positively correlated with the mRNA level (Fig. 3, Supplemental Table S20K-N). Among these loci, two interesting examples were identified: the well-known ripening-associated genes *RIPENING INHIBITOR* (*RIN*, Solyc05g012020, (Vrebalov et al. 2002)) and *FRUITFULL2* (*FUL2*, Solyc03g114830, (Bemer *et al.,* 2012)), which showed higher expression in *phyB1B2* at the IG stage (Supplemental Fig. S5a) and higher sRNA accumulation and sCTGR methylation across their GBs (Supplemental Fig. S5b) compared to WT. The premature expression of these TFs was in agreement with the previously reported anticipation of ripening onset in the *phyB1B2* mutant (Gupta *et al*., 2014). Altogether, these findings revealed: (i) impaired RdDM in BK fruits of *phyB1B2*, indicated by the absence of clusters with less sRNA accumulation (Supplemental Fig. S4); and (ii) that GB RdDM is an important mechanism that positively regulates gene expression in a PHY-mediated manner during fruit development (Fig. 3).

### PHYB1B2-dependent methylation regulates fruit chlorophyll accumulation

The categorization of DEGs associated with differential promoter methylation revealed prominent representation of the photosynthesis category in the fruits of the *phyB1B2* mutant at the IG stage (Fig. 1C). Among the 32 genes, 22 were downregulated and hypermethylated in the promoter region (Supplemental Tables S6 and S17). Most of these genes encode chlorophyll-binding proteins, structural photosystem proteins and chlorophyll biosynthetic enzymes. This might at least partly explain the reduction of 50% in the total chlorophyll level observed in *phyB1B2* IG fruits (Supplemental Fig. S6A). The detailed analysis of the chlorophyll biosynthetic *PROTOCHLOROPHYLLIDE OXIDOREDUCTASE 3* (*POR3,* Solyc07g054210) and two *CHLOROPHYLL A/B BINDING PROTEINs* (*CBP*, Solyc02g070990 and *CAB-3c*, Solyc03g005780) genes showed that their reduced mRNA levels in *phyB1B2* (Supplemental Fig. S6B) correlated with the presence of hypermethylated regions in the promoters (Supplemental Fig. S6C). These results suggest that the transcription of genes involved in chlorophyll metabolism and the photosynthetic machinery in tomato fruits is affected by the PHYB1B2-dependent methylation status of their promoter regions.

### The methylation-mediated regulation of fruit ripening is influenced by PHYB1B2 signalling

In their seminal study, Zhong *et al*. (2013) revealed that the extensive methylation in the promoter regions of ripening-associated genes gradually decreases during fruit development. Interestingly, RNA biosynthesis, which includes transcription factors, was the most abundant functional category among the DEGs that showed DMPs (Fig. 1C). Thus, we examined a set of ripening-associated transcription factors: *RIN*, *NON-RIPENING* (*NOR*, Solyc10g006880 (Mizrahi et al. 1976)), *COLORLESS NORIPENING* (*CNR*, Solyc02g077920, Manning *et al.* (2006)) and *APETALA2a* (*AP2a*, Solyc03g044300, Karlova *et al.* (2011)). The evaluation of the promoter regions clearly showed that while their methylation level decreased from the IG to BK stage in the WT genotype, they remained highly methylated in *phyB1B2* (Fig. 4A). The maintenance of high methylation levels in the promoters of these key regulatory genes at the onset of fruit ripening was highly correlated with their transcriptional downregulation at the BK stage (Fig. 4B).

**Fig. 4.**
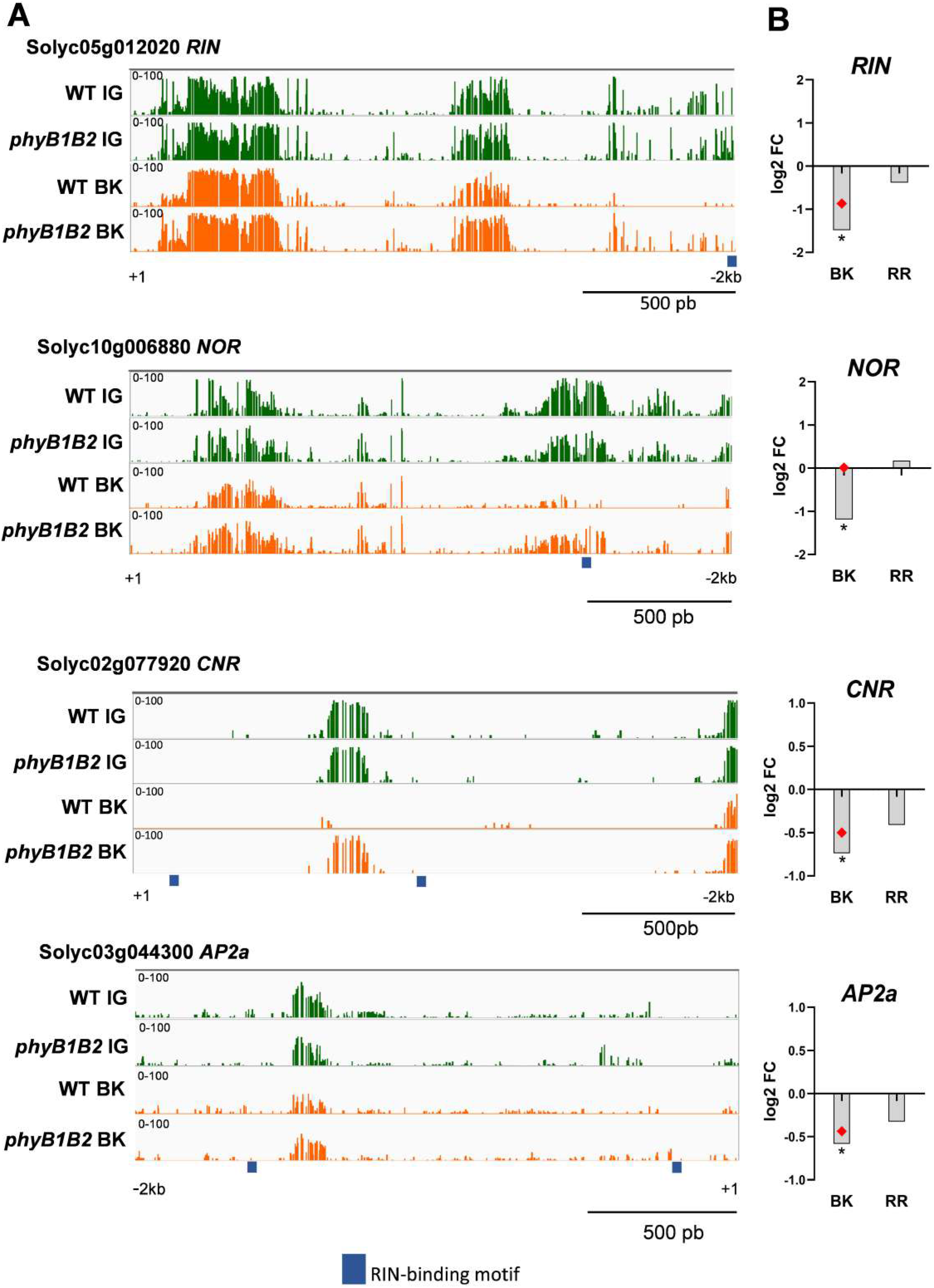
PHYB1/B2 influence on fruit ripening is associated to the promoter demethylation of master ripening-associated transcription factors (A) Differentially methylated promoters of the *RIPENING INHIBITOR* (*RIN*), *NON-RIPENING* (*NOR*), *COLORLESS NORIPENING* (*CNR*) and *APETALA 2a* (*AP2a*) loci between the *phyB1B2* and wild-type (WT) genotypes. Green and orange indicate total mC in immature green (IG) and breaker (BK) fruits, respectively. HY5 and PIF transcription factor binding motifs are denoted with arrows. Thick blue lines indicate RIN binding sites according to ChIP-seq data(Zhong et al. 2013). (B) Relative expression from the RT-qPCR assay of genes encoding master ripening transcription factors in BK and red ripe (RR) fruits from *phyB1B2*. Red dots indicate data from RNA-seq in the same stage. Expression levels represent the mean of at least three biological replicates and are relative to WT. Asterisks indicate statistically significant differences by two-tailed Student’s *t* test compared to WT (* *p*<0.05).

Carotenoid accumulation is probably the most appealing and best investigated trait of tomato fruits; in agreement with previous findings (Bianchetti et al. 2020), ripe *phyB1B2* fruits showed a five-fold reduction in carotenoid content compared to WT (Fig. 5A). With the aim of evaluating whether this effect is a consequence of the methylation-mediated regulation of carotenoid biosynthesis genes, we further analysed the promoters of *PHYTOENE SYNTHASE 1* (*PSY1,* Solyc03g031860), *PHYTOENE DESATURASE* (*PDS,* Solyc03g123760), *15-CIS-ζ-CAROTENE* (*ZISO,* Solyc12g098710) and *ZETA-CAROTENE DESATURASE* (*ZDS,* Solyc01g097810), which, with the exception of *PDS*, were hypermethylated in *phyB1B2* BK fruits (Supplemental Table S11). The mC profile confirmed the presence of hypermethylated regions in all four promoters (Fig. 5B), which might explain the reduced mRNA levels of these genes observed in *phyB1B2* (Fig. 5C).

**Fig. 5.**
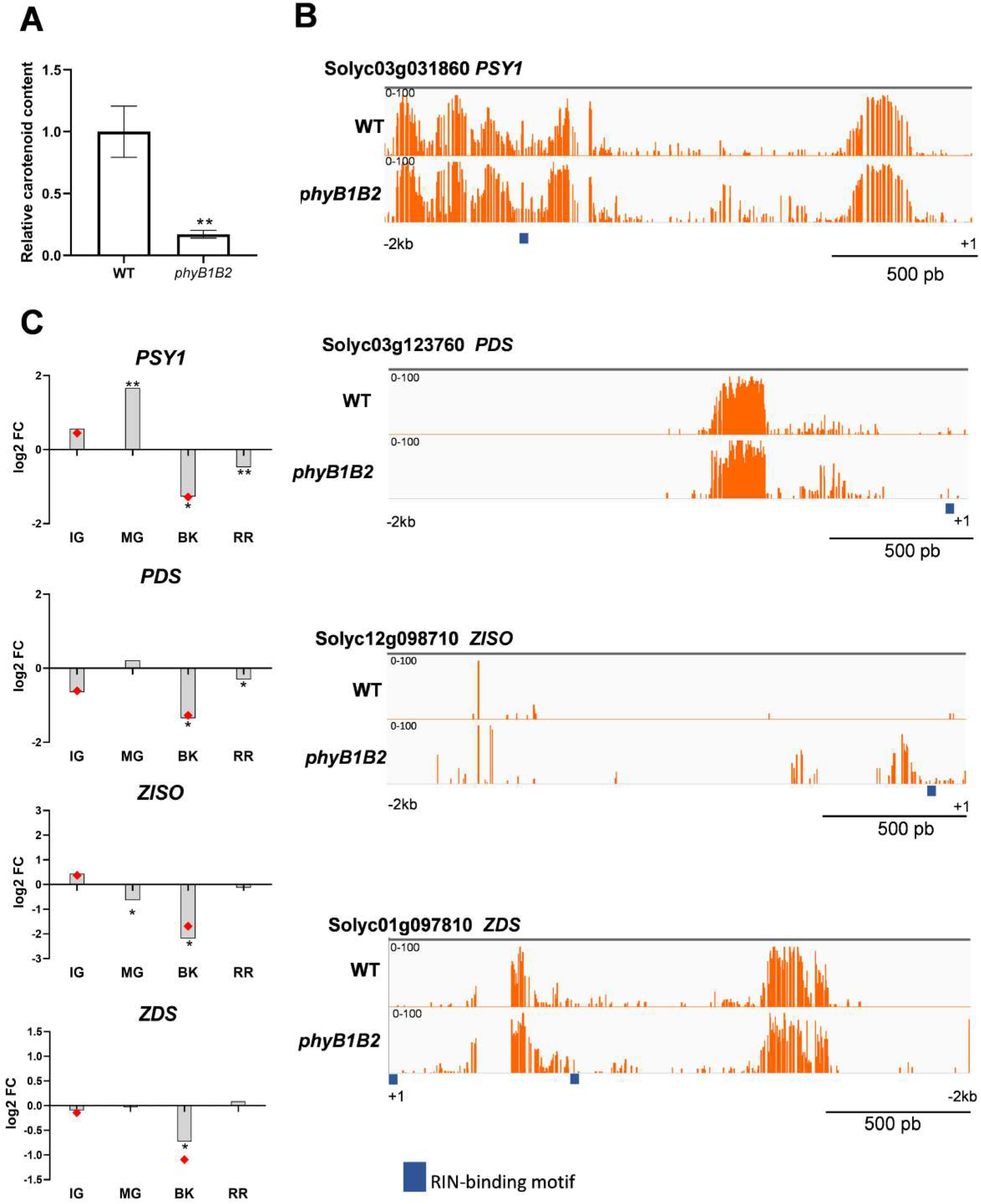
PHYB1/B2-dependent regulation of fruit carotenogenesis relies on the promoter demethylation of key carotenoid biosynthetic genes. (A) Relative contents of total carotenoids in red ripe (RR) fruits from phyB1B2 and wild-type (WT) genotypes. Values represent the mean of at least three biological replicates. Asterisks indicate statistically significant differences by the two-tailed Student’s *t* test between genotypes (** *p*<0.01). (B) Differentially methylated promoter sites of the *PHYTOENE SYNTHASE 1* (*PSY1*), *PHYTOENE DESATURASE* (*PDS*), *15-CIS-ζ-CAROTENE* (*ZISO*) and *ZETA-CAROTENE DESATURASE* (*ZDS*) loci between the *phyB1B2* and WT genotypes. Orange colour indicates total mC in breaker (BK) fruits. Arrows denote HY5 and PIF transcription factor binding motifs. Thick blue lines indicate RIN binding sites according to ChIP-seq data(Zhong et al. 2013). (C) Relative expression of carotenoid biosynthetic enzyme-encoding genes in immature green (IG), mature green (MG), BK and RR fruits from *phyB1B2* determined by RT-qPCR. Red dots indicate data from RNA-seq in the same stage. The expression levels represent the mean of at least three biological replicates and are relative to WT. Asterisks indicate statistically significant differences by the two-tailed Student’s *t* test compared to WT (\**p*<0.05, ** *p*<0.01).

RIN is one of the main TFs controlling ripening-associated genes by directly binding to their promoters. RIN binding occurs in concert with the demethylation of its targets (Zhong et al. 2013). To examine whether RIN binding site methylation could be affected by the *phyB1B2* mutation in the ripening-related master transcription factors and carotenoid biosynthetic gene promoters, we mapped the available RIN ChIP-seq data (Zhong et al. 2013) and performed *de novo* motif discovery (Supplemental Fig. S7). Interestingly, the levels of mCs on the RIN target genes promoters, *NOR*, *CNR* and *AP2a,* were higher in the *phyB1B2* than in WT. Moreover, the *RIN* promoter itself was hypermethylated across the RIN binding site in *phyB1B2* BK fruits, suggesting a positive feedback regulatory mechanism (Fig. 4A). Finally, in the *phyB1B2* mutant, the *PSY1*, *PDS*, *ZISO* and *ZDS* promoters showed higher methylation overlapping with RIN target binding sites (Fig. 5B), indicating that the upregulation of carotenoid biosynthesis genes during tomato ripening is dependent on the PHYB1B2-mediated demethylation of RIN target binding sites. Altogether, our findings showed that PHYB1B2 is a major player in fruit ripening by affecting the promoter demethylation of master transcriptional regulators and carotenoid biosynthesis genes.

### *Cis-*regulatory PIFs/HYx/RIN elements in promoter regions of *phyB1B2* DEGs

The frequency and overrepresentation of PHY-downstream effectors, particularly PIFs and HYx (HY5 and HYH), and RIN binding motifs on *phyB1B2* DEGs promoter regions were evaluated. Three gene datasets were separately analized: *phyB1B2-*upregulated, *phyB1B2*-downregulated and those related to chromatin organization functional category. The proportion of promoters that contains each motif in the analyzed region is depicted in Fig. 6A. After subtracting the background signal, it results evident that the promoter region of the chromatin organization DEGs are overrepresented in PIFs and HYx binding motifs (Fig. 6B). These results suggest that the effect of PHYB1B2 on the expression of the chromatin organization genes is mediated by the downstream effectors: PIFs and HYx. Moreover, RIN binding motif was overrepresented on the three gene datasets evaluated, being higher on the *phyB1B2-*upregulated genes (Fig. 6B).

**Fig. 6.**
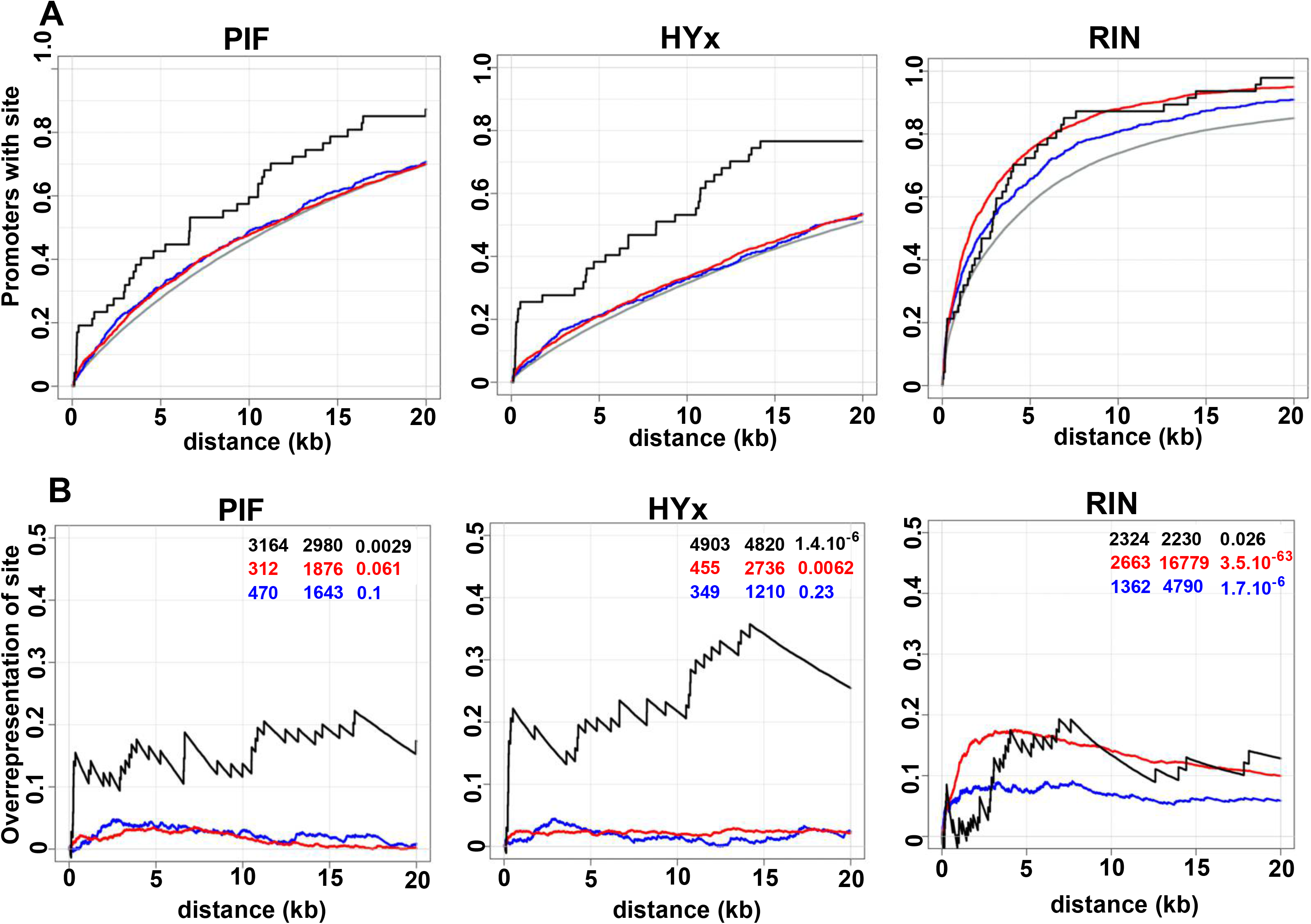
Positional distribution and enrichment of TF binding sites on PHYB1B2 regulated genes. The three gene dataset analysed: upregulated (red), downregulated (blue) and chromatin-remodeling (black) DEGs. (A) Additive gene percentage harbouring the indicated element in comparison with randomly chosen gene set (grey). (B) Over-representation of elements in the regulated genes in comparison to the randomly chosen gene set by subtracting of the curves shown in (A). The enrichment score, z-score and *P*-value for each class of TF are shown from left to right as inset. PIF includes PHYTOCHROME INTERACTING FACTOR 1,3,4,5 and 7 sites; HYx includes LONG HYPOCOTYL 5 (HY5) and HY5 HOMOLOG (HYH) from Jaspar Database.

## DISCUSSION

The dynamic methylation pattern during tomato fruit development has been demonstrated to be a critical ripening regulation mechanism (Zhong et al. 2013; Zuo et al. 2020). DNA demethylation, mainly in the CG context, triggers the activation of genes involved in ripening and is required for pigment accumulation and ethylene synthesis (Zhong et al. 2013; Lang et al. 2017). Simultaneously, the dynamic epigenome during fruit development is strictly regulated by environmental cues (Zhang et al. 2016). The prevailing model establishes PHYs as major components involved in the coordination of fruit physiology with the ever-changing light and temperature environmental conditions (Alves et al. 2020; Bianchetti et al. 2020). Thus, we explored the link between fruit epigenome reprogramming and these well-established light and temperature sensors (Legris *et al*., 2016).

Our data clearly showed that *phyA* and *phyB1B2* deficiencies modified the epigenome profile through methylome and sRNAome reprogramming. In particular, PHY-mediated DMPs and GB methylation were associated to transcriptome alterations that affected tomato fruit development; thus, indicating that active PHYs regulate, at least in part, the ripening-associated demethylation previously reported (Zhong *et al*., 2013). However, the massive alteration of methylation patterns observed in *phy* mutants suggests the existence of a still unclear genome regulatory mechanism.

The *phyA* and *phyB1B2* mutants showed a positive correlation between cluster sRNA accumulation, target methylation in GB and mRNA levels. In angiosperms, GB methylation has been associated with constitutively expressed genes (Bewick and Schmitz 2017; Lu et al. 2015); however, PHY deficiency intriguingly seems to deregulate this mechanism, affecting the temporally expression of regulated genes. The *RIN* and *FUL2* examples analysed here clearly showed that sRNA accumulation and methylation were mainly located near transposable elements (TEs) (Supplemental Fig. S5). It is known that the insertion of TEs within GB can disrupt gene expression; thus, methylation-mediated TE silencing and GB methylation are evolutionarily linked (Bewick and Schmitz 2017). The enhancement of TE-associated DNA methylation in GB (Fig. 3C) and the absence of clusters with less sRNA accumulation in BK compared to the IG stage in *phyB1B2* (Supplemental Fig. S4) might be explained by the overexpression of canonical RdDM genes: Solyc12g008420 and Solyc06g050510 encode homologs of RNA-DEPENDENT RNA POLYMERASE (RDRP) and the associated factor SNF2 DOMAIN-CONTAINING PROTEIN CLASSY 1 (CLSY1), respectively, both of which were upregulated in BK fruits from *phyB1B2* plants (Table S4). Similarly, Solyc09g082480 and Solyc03g083170, which were also upregulated in *phyB1B2* BK fruits, are homologs of *A. thaliana* RNA-DIRECTED DNA METHYLATION 1 (RDM1) and DEFECTIVE IN MERISTEM SILENCING 3 (DMS3), respectively. The protein products of these genes, together with DEFECTIVE IN RNA-DIRECTED DNA METHYLATION 1 (DRD1), form the DDR complex, which enables RNA Pol V transcription (Pikaard and Scheid 2014). To our knowledge, this is the first report to associate PHY-mediated sRNA accumulation and DNA methylation with mRNA levels in plants.

Several pieces of evidence have shown that PHYB1B2 has a more substantial impact on tomato epigenome regulation than PHYA. For example, BK fruits from *phyB1B2* displayed (i) a large number of DEGs associated with chromatin organization (Fig. 1C); (ii) overall promoter hypermethylation in the CG context (Fig. 2B); (iii) the highest number of DEGs associated with DMPs (Supplemental Fig. S2); and (iv) half the number of DMPs associated with DEGs between the IG and BK stages compared to the WT (Supplemental Fig. S3). In order to understand how *phyB1B2* mutation resulted in this massive epigenomic alteration, we closely looked at the DEGs related to chromatin organization functional category.

The chromomethylase *SlMET1L* (Solyc01g006100) (also referred to as *SlCMT3* (Gallusci et al. 2016) displays the highest transcript abundance in immature fruits, which declines towards the fully ripe stage (Cao et al. 2014). In line with the higher level of DNA methylation, our transcriptome analysis showed that *SlMET1L* was upregulated in *phyB1B2* BK fruits. Conversely, *SlROS1L* demethylase (Solyc09g009080, Cao *et al.*, 2014), also referred as *SlDML1* (Liu et al. 2015), was also upregulated in *phyB1B2* BK fruits. Although, it might seem contradictory at first glance, it has been reported that the *Arabidopsis thaliana ROS1* gene promoter contains a DNA methylation monitoring sequence (MEMS) associated with a Helitron transposon, which is methylated by AtMET1, positively regulating *AtROS1* gene expression (Lei et al. 2015). Similarly, *SlROS1L* harbours two transposable elements within its promoter and showed a higher methylation level in *phyB1B2* than in the WT genotype, suggesting a similar regulatory mechanism in tomato (Supplemental Fig. S8, Supplemental Table S15).

The tomato homolog of *A. thaliana* DECREASED DNA METHYLATION 1 (DDM1, Solyc02g085390) showed higher mRNA expression in *phyB1B2* mutant BK fruits than in their WT counterparts. DDM1 is a chromatin remodelling protein required for maintaining DNA methylation in the symmetric cytosine sequence (Zemach et al. 2013), which can be associated with the CG context hypermethylation observed in *phyB1B2* (Fig. 2A).

Several histone modifiers showed altered expression in BK fruits from the *phyB1B2* mutant (Supplemental Table S8). The methylation of lysine residues 9 and 27 on H3 is associated with repressed genes. Histone lysine methyltransferases are classified into five groups based on their domain architecture and/or differences in enzymatic activity (Pontvianne et al. 2010). The BK fruits of the *phyB1B2* mutant displayed three differentially expressed lysine methyltransferases: Solyc03g082860, an upregulated H3K27 Class IV homolog; and two H3K9 Class V homologs, Solyc06g008130 and Solyc06g083760, showing lower and higher expression than WT fruits, respectively. Histone arginine methylation is catalysed by a family of enzymes known as protein arginine methyltransferases (PRMTs). Solyc12g099560, a PRMT4a/b homolog, was upregulated in *phyB1B2* BK fruits. Interestingly, in *A. thaliana*, PRMT4s modulate key regulatory genes associated with the light response (Hernando et al. 2015), reinforcing the link between the PHYB1B2 photoreceptors and epigenetic control. Finally, tomato histone demethylases have been recently identified. *SlJMJ6*, whose expression peaks immediately after the BK stage, has been characterized as a positive regulator of fruit ripening by removing the H3K27 methylation of ripening-related genes, and *SlJMJ6*-overexpressing lines show increased carotenoid levels (Li et al. 2020). *SlJMJC1* (Solyc01g006680), which exhibits the same expression pattern (Li et al. 2020), is downregulated in the *phyB1B2* mutant, suggesting that this gene might exhibit similar regulatory function to its paralog, inducing ripening in a PHYB1B2-dependent manner (Figs. 4 and 5).

Histone deacetylation plays a crucial role in the regulation of eukaryotic gene activity and is associated with inactive chromatin (Zhang et al. 2018). Histone deacetylation is catalysed by histone deacetylases (HDACs). Fifteen HDACs were identified in the tomato genome (Zhao et al. 2015). Among these, *SlHDA10* (Solyc01g009120) and *SlHDT3* (Solyc11g066840) were found to be downregulated and upregulated in *phyB1B2* BK fruits, respectively. SlHDA10 is localized in the chloroplast, and its transcript is highly expressed in photosynthetic tissues (Zhao et al. 2015); whether SlHDA10 deacetylates chloroplast proteins by silencing photosynthesis-related genes remains to be determined. Although *SlHDT3* is mainly expressed in immature stages of fruit development and its expression declines with ripening, its silencing results in delayed ripening and reduced *RIN* expression and carotenogenesis. On the other hand, the expression level of *SlHDT3* is increased in ripening-deficient mutants such as *Nr* or *rin* (Guo et al. 2017). Our results showed that *phyB1B2* mutant fruits displayed higher expression of *SlHDT3* and reduced *RIN* transcript levels at the BK stage, suggesting reciprocal regulation between these two factors. Thus, we propose that during the IG stage, *SlHDT3* is highly expressed, contributing to the epigenetic inhibition of ripening. The reduction in *SlHDT3* expression towards BK releases DNA methylation that, in turn upregulates *RIN* tunning ripening-related epigenetic reprogramming and contributing to explain the high methylation levels observed in the *phyB1B2* mutant (Fig. 2).

Fruit ripening is a key trait for fitness and several alternative regulatory mechanisms guarantee the success of this process. This is most probably the reason why a single initiating signal has not been identified (Giovannoni et al. 2017). A complex interactive module involving DNA methylation level and tomato ripening- transcription factors was described (Zhong et al. 2013; Zuo et al. 2020). On the other hand, the link between chromatin remodelling and light signalling has been previously reported (Fisher and Franklin, 2011). Here, the comprehensive analysis of the experimental evidences allowed us to propose that PHYs, specially PHYB1B2, are important factors that participates in the crosstalk among chromatin organization and transcriptional regulators. The enrichment of PIF and HYx *cis*-regulatory motifs among the promoters of *phyB1B2-*DEGs associated with chromatin organization suggests that these PHY downstream factors regulate these genes that, in turn, trigger ripening-associated DNA demethylation. Epigenome reprogramming results in the adjustment of transcriptome including the induction of RIN master TF. The enrichment of hypermethylated RIN binding sites on the promoters of key ripening TFs (CNR, NOR and AP2a), including RIN itself, in *phyB1B2*, indicates their RIN-mediated induction. These observations together with the fact that *rin* mutant is impaired in ripening-associated demethylation (Zhong et al., 2013), allow us to propose a positive regulatory loop between PHYs downstream effectors-and RIN-mediated DNA demethylation, driving the transcriptional regulation of ripening associated TFs and, finally, to a shift in the expression profile along fruit development (Fig 7). The vast reservoir of data released here brings a new level of understanding about how epigenetic mechanisms orchestrate the response to PHY-mediated light and temperature fluctuations affecting important agronomical traits in fleshy fruits.

**Fig. 7.**
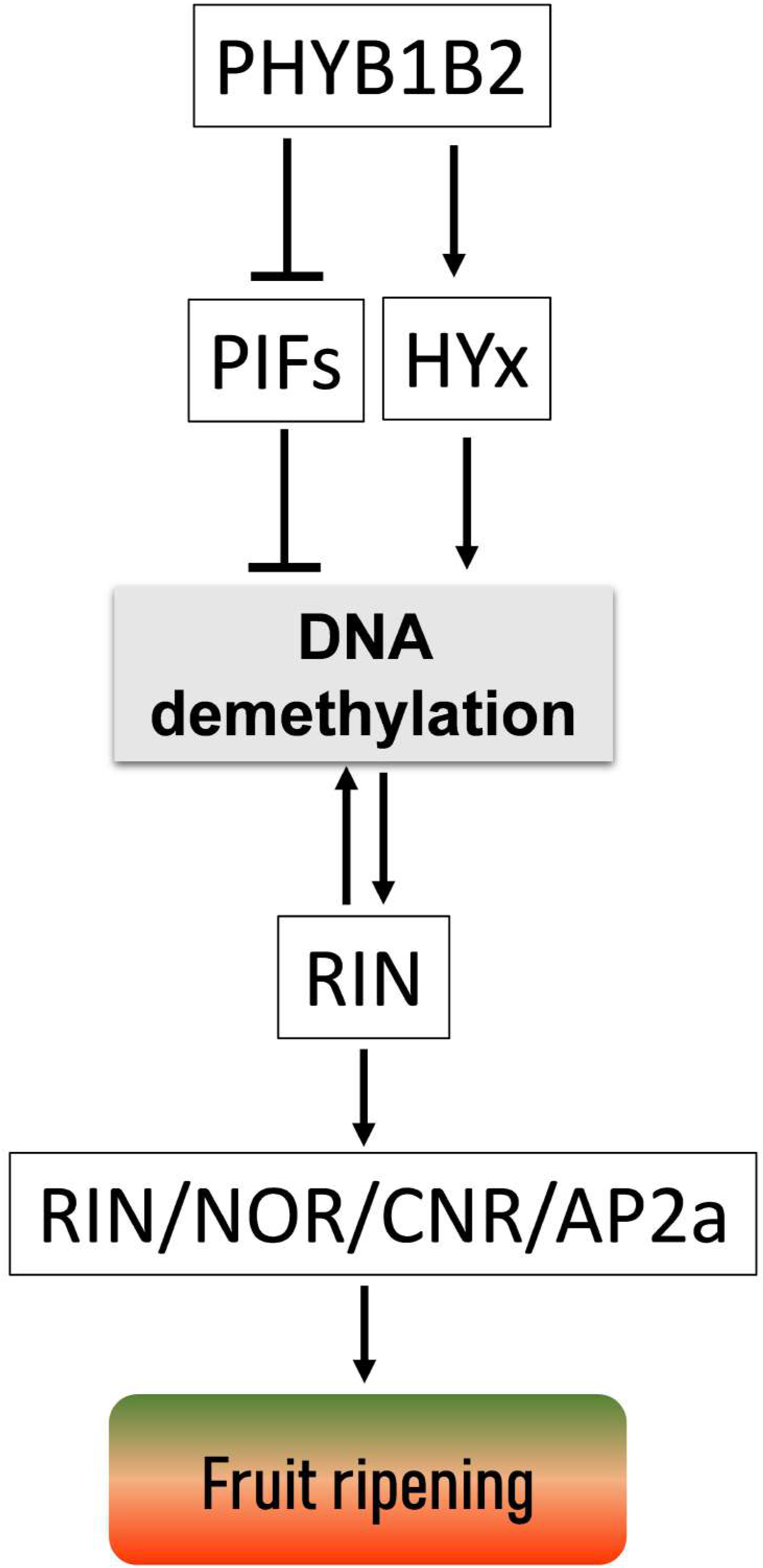
Conceptual model linking PHYB1B2 receptors, epigenetic mechanisms of gene expression regulation and fruit ripening. Active PHYB1B2, through the inactivation of PIFs and HYx stabilization, regulate the expression of chromatin organization associated genes resulting in DNA demethylation and the expression induction of RIN ripening master TF. RIN targets include chromatin organization genes resulting in a positive feedback loop. Moreover, RIN enhances its own transcription, as well as other TFs (such as NOR, CNR and AP2a) that finally induce a myriad of effectors triggering ripening.

## METHODS

### Plant material, growth conditions and sampling

*phyA* single and *phyB1B2* double mutants in the *Solanum lycopersicum* (cv. MoneyMaker) genetic background were previously characterized (Kerckhoffs et al. 1996; Lazarova et al. 1998; Kerckhoffs et al. 1999). Plants were grown in a glasshouse at the Instituto de Biociências, Universidade de São Paulo, 23°33′55′′S 46°43′51′′W. Tomato seeds were grown in 9L pots containing a 1:1 mixture of commercial substrate and expanded vermiculite, supplemented with 1 g L^−1^ of NPK 10:10:10, 4 g L^−1^ of dolomite limestone (MgCO_3_ + CaCO_3_) and 2 g L^−1^ thermophosphate at 24/18 °C under a 16/8 h light/dark cycle under 230-250 μmol photons m^−2^ s^−1^ irradiation and a relative humidity of 55%. Since it is known that PHYs can affect ripening time (Gupta *et al*., 2014), fruits were sampled at the same development stage instead of necessary the same age. Five replicates per genotype were cultivated, and fruits were sampled at the immature green (15 mm diameter), mature green (when the placenta displays a jelly aspect), breaker (beginning of ripening process when the fruit shows the first yellowish colouration) and red ripe (7 days after the breaker stage) stages. All fruits were harvested at the same time of day with four biological replicates (each replicate was composed of a single fruit per plant). The columella, placenta, and seeds were immediately removed, and the remaining tissues were frozen in liquid nitrogen, ground and freeze-dried for subsequent analysis.

### Transcriptional profile

Total RNA was extracted from immature green and breaker stage fruits with three independent biological replicates of each genotype using a Promega ReliaPrep RNA tissue kit according to the manufacturer’s instructions. The RNA concentration was determined with a spectrophotometer (Nanodrop ND-1000; NanoDrop Technologies, Wilmington, DE, U.S.A.), RNA quality was assessed with a BioAnalyzer 2100 (Agilent Technologies), and RNA libraries were constructed following the recommendations of an Illumina Kit (Directional mRNA-Seq Sample Preparation) and sequenced using the Illumina NovaSeq 6000 System. Each library was sequenced, generating approximately 20 million 150 bp paired end reads per sample. The raw sequencing reads that were generated were analysed with FastQC (http://www.bioinformatics.babraham.ac.uk/projects/fastqc/) and were filtered and cleaned using Trimmomatic (Bolger et al. 2014) (Parameters: ILLUMINACLIP: TruSeq3-PE.fa:2:30:10LEADING:3 TRAILING:3 SLIDINGWINDOW:4:20 MINLEN:50). At least 95% (19.1-27.9 M) of the reads met the quality criteria and were mapped to the tomato reference genome sequence SL3.0 with the ITAG3.2 annotation using STAR v2.4.2. allowing one mismatch (Dobin et al. 2013), approximately 84% of the reads were uniquely mapped (Supplemental Table S1) and were used for statistical analysis.

### Reverse transcription quantitative PCR (RT-qPCR)

Total RNA extraction was performed with the ReliaPrep™ RNA Cell and Tissue Miniprep System (Promega), and cDNA synthesis was conducted with SuperScript™ IV Reverse Transcriptase (Invitrogen). The primers used for qPCR are listed in Supplemental Table S21. RT-qPCR was performed in a QStudio6 – A1769 PCR Real-Time thermocycler using 2X Power SYBR Green Master Mix in a final volume of 10 μL. Absolute fluorescence data were analysed using LinRegPCR software to obtain Ct and primer efficiency values. Relative mRNA abundance was calculated and normalized according to the ΔΔCt method using *EXPRESSED* and *CAC* as reference genes (Expósito-Rodríguez et al. 2008).

### MethylC-Seq analysis

gDNA (~5 g) was extracted from a pool of the same three biological replicates used in the transcriptome analyses, obtained from three IG and BK fruit samples per genotype, using the DNeasy Plant maxi kit (Qiagen). The libraries were prepared with the EZ DNA Methylation-Gold Kit (Zymo Research) and the Accel-NGS^^®^^ Methyl-Seq DNA Library Kit (Swift Biosciences) and further sequenced using the Illumina NovaSeq 6000 platform. Over 240 M reads were sequenced from each genotype and stage. Raw reads were screened for quality using Trimmomatic (Bolger et al. 2014) (parameters: ILLUMINACLIP:TruSeq3-PE.fa:2:30:10 LEADING:3 TRAILING:3 SLIDINGWINDOW:4:20 MINLEN:50). Mapping to the tomato reference genome sequence SL3.0 and the assessment of global methylation status were performed using Bismark (Krueger and Andrews 2011) (parameters: bismark -q --bowtie2 --non_directional -N 1 -p 4), and the methylation status of DNA in the three possible contexts (CG, CHG and CHH) was distinguished. At least 130 M reads were uniquely mapped (Supplemental Table S9). The Bioconductor package methylKit (Akalin et al. 2012) was used for the detection of methylation levels across the analysed regions: promoters (2 kb upstream of transcription start site) and sRNA cluster-targeted genome regions (sCTGRs). Only Cs with 10X coverage were considered. Methylation differences with a FDR < 0.05 in each comparison (WT vs *phyA*; WT vs *phyB1B2*) were recorded as differentially methylated promoters (DMPs) or differentially methylated sCTGRs. Differential methylation in the CG, CHG and/or CHH context was considered if the region contained, at least, 10 differentially methylated Cs in the corresponding context. Finally, for the comparison of global methylation levels between genotypes, only common Cs with at least 10X coverage in all samples were analysed.

### sRNAome profile

sRNA extraction and quality parameters were determined from the same replicates described above in the “Transcriptional profile” section. After RNA integrity confirmation, libraries were prepared using a TruSeq Small RNA Library Prep and sequenced using the Illumina HiSeq 4000 platform to generate a read length of 50 bp. The raw sequencing reads that were generated were quality trimmed with Trimmomatic (Bolger et al. 2014) to retain reads of 18-24 nt in length (parameters: ILLUMINACLIP:TruSeq3-SE:2:30:10 LEADING:3 TRAILING:3 SLIDINGWINDOW:4:15 MINLEN:18 AVGQUAL:25). A minimum of 38% (WT/breaker/A) and a maximum of 85% (WT/immature green/A) of the reads achieved the quality criteria and were used for further analyses (Supplemental Table S20A). All libraries were aligned to genome version SL3.0 using ShortStack v3.8.1(Axtell 2013) with default parameters (allowing the distribution of multimapping reads according to the local genomic context). Then, the *de novo* identification of clusters of sRNAs was performed for all libraries, and individual counts for each library and cluster were obtained using the same software.

### Statistical analysis for RNAseq and sRNAome

Genes/sRNA clusters with read/count numbers smaller than two per million were removed. Read/count values were normalized according to the library size factors. Statistical analyses were performed with edgeR from Bioconductor^®^ (Robinson et al. 2009; McCarthy et al. 2012) using a genewise negative binomial generalized linear model with the quasi-likelihood test (Chen et al. 2016) and a cutoff of the false discovery rate (FDR) ≤ 0.05.

### Gene functional categorization

The DEGs were functionally categorized with MapMan application software (Thimm et al. 2004) followed by hand-curated annotation using MapMan categories.

### *In silico* regulatory motif predictions and RIN ChIP-seq analyses

RIN ChIPseq reads were downloaded from the Sequence Read Archive (SRA) (accession SRX15083 (Zhong et al. 2013), mapped to tomato genome version SL3.0 with STAR (Dobin et al. 2013) (version 2.7.3X, parameters: outFilterMismatchNmax 3, alignEndsType EndToEnd, alignIntronMax 5), and peak calling was performed using Macs2 (Zhang et al. 2008) (version 2.2.7.1, default parameters). Regions of 200 bp centred on the top-scoring peaks (score>100, n=327) were retrieved and the binding motif was inferred *de novo* by using the MEME algorithm (Bailey et al. 2015).

In order to analyse the relative abundance of light regulation associated *cis*-elements, their position frequency matrices (PFM) were retrieve from JASPAR 2020 database (Fornes et al. 2019) for PIFs and HYx (HY5 and HYH) and; from the peak calling of ChIPseq data for RIN (Zhong et al., 2013). The PFMs were scanned with Fimo (Bailey et al. 2015), *P*-value < 1e^−5^ along SL3.0 genome. A 20 Kb region upstream the transcription start site (TSS) was examined for the presence of the TFBSs (Transcriptional Factor Binding Sites). The association was calculated from the accumulative number of genes harbouring a determined *cis*-regulatory element in a specific set of regulated genes, against whole genome random expectation. The signal to noise ratio for each position was calculated as the enrichment score (ES) substracting the regulated genes-set to all annotated promoters. Later, an associated z-score and *P*-value for each class of TF were obtained from the ES distribution of 1000 random samples set.

### Carotenoid and chlorophyll analysis

Chlorophyll, phytoene, phytofluene, lycopene, β-carotene and lutein levels were extracted and determined via HPLC with a photodiode array detector as previously described by Lira *et al.* (2017).

### Statistical analysis of RT-qPCR and metabolites

Statistical analyses of the RT-qPCR (Student’s t-test, *p* ≤ 0,05) and metabolic data (ANOVA, Tukey’s test. *p* ≤ 0,05) were performed with InfoStat/F software (http://www.infostat.com.ar).

## DATA ACCESS

All high-throughput sequencing data reported in this paper have been submitted to the Sequence Read Archive (SRA) under NCBI Bioproject PRJNA646733, with accession numbers SUB7763724, SUB7782168 and SUB7791358 for RNAseq, WGBS and small RNAseq, respectively.

## ACKNOWLEDGMENTS

This work was supported by FAPESP (Fundação de Amparo à Pesquisa do Estado de São Paulo, Grant Number #2016/01128-9); RB was a recipient of a FAPESP fellowship (#2017/24354-7). MR was a recipient of a CNPq fellowship. LB was a recipient of a CAPES-PRINT scholarship (88887.370243/2019-00).

## AUTHOR CONTRIBUTIONS

RB performed most of the experiments and analysed the data; LB, NB, LAH and RZ analysed the data; DR performed the experiments; RB, LF, MR and LB conceived the project, designed the experiments and wrote the paper, which was revised and approved by all authors. LB agrees to serve as the author responsible for contact and ensures communication.

## Supplementary Figures

**Fig. S1.** Global methylation status in *phyA*, *phyB1B2* and WT at IG and BK stage for mCG (A), mCHG (B) and mCHH (C) contexts. Density plot ofgenes, transposable elements (TEs) and sRNAs clusters were estimated by the number of nucleotides covered per million. Methylation levels for CG, CHG and CHHcontexts, are 40-90%; 25-80% and 10-30%, respectively. Cytosine density for CG, CHG and CHH contexts, are 0 - 17,488; 0 - 13,105 and 0 - 103,903 per million,respectively.

**Fig. S2.** mRNA level alterations associated with differences in the promoter methylation in*phyA* and *phyB1B2* compared to WT. Scatter plots show the DEGs that showed DMPs at immature green (IG) and breaker (BK) fruit stages, the axes indicate the variation between both parameters compared to WT(DEGs, FDR < 0.05; DMPs, FDR < 0.05) in CG, CHG and CHH contexts. Only DMPs with changes inmethylation levels > 5% are shown.

**Fig. S3.** Comparison of the association of DMPs and DEGs between immature green and breaker stages within each genotype. DMPs: differentially methylated promoters; DEGs: differentially expressed genes.

**Fig. S4.** Methylation levels in sRNA cluster targeted genomic region (sCTGR) and small RNA accumulation changes between IG and BK stages in WT, *phyA* and *phyB1B2* genotypes. Scatter plots show the relationship between the small RNA accumulation and differential methylation on their target genomic regions between immature green (IG) and breaker (BK) fruit stages.

**Fig. S5.** Methylation across promoter and gene body regions differentially affects gene expression. (A) Relative expression of *RIPENING INHIBITOR* (*RIN*) and *FRUITFULL 2* (*FUL2*) in immature green (IG) and mature green (MG) fruits from *phyB1B2* determined by RT-qPCR. Red dots indicate data from RNA-seq in the same stage. Expression levels represent the mean of at least three biological replicates and are relative to the wild type (WT). Asterisks indicate statistically significant differences by the two-tailed Student′s *t* test compared to WT (* *p*<0.05). (B) Differential gene body methylation (green bars) and sRNA accumulation (black bars) within *RIN* and *FUL2* in IG fruits from the *phyB1B2* and WT genotypes.

**Fig. S6.** PHYB1/B2-dependent methylation regulates fruit chlorophyll. (A) Relative content of total chlorophyll in IG fruits from *phyB1B2* and WT genotypes. Values represent the mean of at least three biological replicates. Asterisks indicate statistically significant differences by the two-tailed Student’s *t* test between genotypes (** *p*<0.01). (B) Relative expression of *CHLOROPHYLL A/B BINDING PROTEINs* (*CBP* and *CAB-3c*) and *PROTOCHLOROPHYLLIDE OXIDOREDUCTASE 3* (*POR3*) in IG fruits from *phyB1B2* determined by RNA-seq. Expression levels represent the mean of at least three biological replicates and are relative to WT. Asterisks indicate statistically significant differences compared to WT (* FDR≤0.05). (C) Differential promoter methylation in *CBP*, *CAB-3c* and *POR3* in IG fruits from the *phyB1B2* and WT genotypes. HY5 and PIF transcription factor binding motifs are denoted with arrows.

**Fig. S7.** RIN motif *de novo* discovered using MEME algorithm. Consensus sequence CCWWWWWWGG (CC(6W)GG) and extended TTWCCWWWWWWGGWAA length=16.

**Fig. S8.** DMP sites in the *ROS1L* locus in WT and *phyB1B2* BK fruits. Red rectangles indicate the presence of transposable element (TE).

## REFERENCES

Akalin A, Kormaksson M, Li S, Garrett-Bakelman FE, Figueroa ME, Melnick A, Mason CE. 2012. MethylKit: a comprehensive R package for the analysis of genome-wide DNA methylation profiles. Genome Biol 13: R87. http://genomebiology.com/2012/13/10/R87.

Alves FRR, Lira BS, Pikart FC, Monteiro SS, Furlan CM, Purgatto E, Pascoal GB, Andrade SC da S, Demarco D, Rossi M, et al. 2020. Beyond the limits of photoperception: constitutively active PHYTOCHROME B2 overexpression as a means of improving fruit nutritional quality in tomato. Plant Biotechnol J n/a. https://doi.org/10.1111/pbi.13362.

Axtell MJ. 2013. ShortStack: Comprehensive annotation and quantification of small RNA genes. Rna 19: 740–751.

Bailey TL, Johnson J, Grant CE, Noble WS. 2015. The MEME Suite. Nucleic Acids Res 43: W39–W49. https://pubmed.ncbi.nlm.nih.gov/25953851.

Bemer M, Karlova R, Ballester AR, Tikunov YM, Bovy AG, Wolters-Arts M, Rossetto P de B, Angenent GC, de Maagd RA. 2012. The Tomato FRUITFULL Homologs TDR4/FUL1 and MBP7/FUL2 Regulate Ethylene-Independent Aspects of Fruit Ripening. Plant Cell 24: 4437 LP – 4451. http://www.plantcell.org/content/24/11/4437.abstract.

Bertrand C, Benhamed M, Li Y-F, Ayadi M, Lemonnier G, Renou J-P, Delarue M, Zhou D-X,. 2005. Arabidopsis HAF2 gene encoding TATA-binding protein (TBP)-associated factor TAF1, is required to integrate light signals to regulate gene expression and growth. J Biol Chem 280: 1465–1473.

Bewick AJ, Schmitz RJ. 2017. Gene body DNA methylation in plants. Curr Opin Plant Biol 36: 103–110. http://www.sciencedirect.com/science/article/pii/S1369526616301297.

Bianchetti R, De Luca B, de Haro LA, Rosado D, Demarco D, Conte M, Bermudez L, Freschi L, Fernie AR, Michaelson L V, et al. 2020. Phytochrome-Dependent Temperature Perception Modulates Isoprenoid Metabolism. Plant Physiol 183: 869 LP – 882. http://www.plantphysiol.org/content/183/3/869.abstract.

Bolger AM, Lohse M, Usadel B. 2014. Trimmomatic: A flexible trimmer for Illumina sequence data. Bioinformatics 30: 2114–2120.

Bourbousse C, Barneche F, Laloi C. 2020. Plant Chromatin Catches the Sun. Front Plant Sci 10: 1728. https://www.frontiersin.org/article/10.3389/fpls.2019.01728.

Cao D, Ju Z, Gao C, Mei X, Fu D, Zhu H, Luo Y, Zhu B. 2014. Genome-wide identification of cytosine-5 DNA methyltransferases and demethylases in Solanum lycopersicum. Gene 550: 230–237. http://www.sciencedirect.com/science/article/pii/S0378111914009603.

Carlson KD, Bhogale S, Anderson D, Tomanek L, Madlung A. 2019. Phytochrome a regulates carbon flux in dark grown tomato seedlings. Front Plant Sci 10: 1–20.

Chen Y, Lun ATL, Smyth GK. 2016. From reads to genes to pathways: differential expression analysis of RNA-Seq experiments using Rsubread and the edgeR quasi-likelihood pipeline. F1000Research 5: 1438. https://www.ncbi.nlm.nih.gov/pubmed/27508061.

Cheng Y-L, Tu S-L,. 2018. Alternative Splicing and Cross-Talk with Light Signaling. Plant Cell Physiol 59: 1104–1110. https://doi.org/10.1093/pcp/pcy089.

Cho J-N, Ryu J-Y, Jeong Y-M, Park J, Song J-J, Amasino RM, Noh B, Noh Y-S,. 2012. Control of seed germination by light-induced histone arginine demethylation activity. Dev Cell 22: 736–748.

Chua YL, Brown AP, Gray JC. 2001. Targeted histone acetylation and altered nuclease accessibility over short regions of the pea plastocyanin gene. Plant Cell 13: 599–612.

Chua YL, Watson LA, Gray JC. 2003. The transcriptional enhancer of the pea plastocyanin gene associates with the nuclear matrix and regulates gene expression through histone acetylation. Plant Cell 15: 1468–1479.

Corem S, Doron-Faigenboim A, jouffroy O, Maumus F, Arazi T, Bouché N. 2018. Redistribution of CHH Methylation and Small Interfering RNAs across the Genome of Tomato ddm1 Mutants. Plant Cell tpc.00167.2018.

Ding CJ, Liang LX, Diao S, Su XH, Zhang BY. 2018. Genome-wide analysis of day/night DNA methylation differences in Populus nigra. PLoS One 13: 1–14.

Ding J, Shen J, Mao H, Xie W, Li X, Zhang Q. 2012. RNA-directed DNA methylation is involved in regulating photoperiod-sensitive male sterility in rice. Mol Plant 5: 1210–1216.

Dobin A, Davis CA, Schlesinger F, Drenkow J, Zaleski C, Jha S, Batut P, Chaisson M, Gingeras TR. 2013. STAR: ultrafast universal RNA-seq aligner. Bioinformatics 29: 15–21. https://www.ncbi.nlm.nih.gov/pubmed/23104886.

Ernesto Bianchetti R, Silvestre Lira B, Santos Monteiro S, Demarco D, Purgatto E, Rothan C, Rossi M, Freschi L. 2018. Fruit-localized phytochromes regulate plastid biogenesis, starch synthesis, and carotenoid metabolism in tomato. J Exp Bot 69: 3573–3586. https://doi.org/10.1093/jxb/ery145.

Expósito-Rodríguez M, Borges AA, Borges-Pérez A, Pérez JA. 2008. Selection of internal control genes for quantitative real-time RT-PCR studies during tomato development process. BMC Plant Biol 8: 131. http://www.pubmedcentral.nih.gov/articlerender.fcgi?artid=2629474tool=pmcentrezrendertype=abstract.

Fornes O, Castro-Mondragon JA, Khan A, van der Lee R, Zhang X, Richmond PA, Modi BP, Correard S, Gheorghe M, Baranašić D, et al. 2019. JASPAR 2020: update of the open-access database of transcription factor binding profiles. Nucleic Acids Res 48: D87–D92. https://doi.org/10.1093/nar/gkz1001.

Gallusci P, Hodgman C, Teyssier E, Seymour GB. 2016. DNA Methylation and Chromatin Regulation during Fleshy Fruit Development and Ripening. Front Plant Sci 7: 807. https://www.frontiersin.org/article/10.3389/fpls.2016.00807.

Galvão VC, Fankhauser C. 2015. Sensing the light environment in plants: photoreceptors and early signaling steps. Curr Opin Neurobiol 34: 46–53.

Giovannoni J, Nguyen C, Ampofo B, Zhong S, Fei Z. 2017. The Epigenome and Transcriptional Dynamics of Fruit Ripening. Annu Rev Plant Biol 68: 61–84.

Gramegna G, Rosado D, Sánchez Carranza AP, Cruz AB, Simon-Moya M, Llorente B, Rodríguez-Concepcíon M, Freschi L, Rossi M. 2019. PHYTOCHROME-INTERACTING FACTOR 3 mediates light-dependent induction of tocopherol biosynthesis during tomato fruit ripening. Plant Cell Environ 42: 1328–1339. https://doi.org/10.1111/pce.13467.

Guo JE, Hu Z, Li F, Zhang L, Yu X, Tang B, Chen G. 2017. Silencing of histone deacetylase SlHDT3 delays fruit ripening and suppresses carotenoid accumulation in tomato. Plant Sci 265: 29–38. https://doi.org/10.1016/j.plantsci.2017.09.013.

Gupta SK, Sharma S, Santisree P, Kilambi HV, Appenroth K, Sreelakshmi Y, Sharma R. 2014. Complex and shifting interactions of phytochromes regulate fruit development in tomato. Plant Cell Environ 37: 1688–1702. https://doi.org/10.1111/pce.12279.

Hernando CE, Sanchez SE, Mancini E, Yanovsky MJ. 2015. Genome wide comparative analysis of the effects of PRMT5 and PRMT4/CARM1 arginine methyltransferases on the Arabidopsis thaliana transcriptome. BMC Genomics 16: 192. https://doi.org/10.1186/s12864-015-1399-2.

Ibarra SE, Auge G, Sánchez RA, Botto JF. 2013. Transcriptional programs related to phytochrome a function in arabidopsis seed germination. Mol Plant 6: 1261–1273.

Kaiserli E, Perrella G, Davidson ML. 2018. Light and temperature shape nuclear architecture and gene expression. Curr Opin Plant Biol 45: 103–111. http://dx.doi.org/10.1016/j.pbi.2018.05.018.

Karlova R, Rosin FM, Busscher-Lange J, Parapunova V, Do PT, Fernie AR, Fraser PD, Baxter C, Angenent GC, de Maagd RA. 2011. Transcriptome and Metabolite Profiling Show That APETALA2a Is a Major Regulator of Tomato Fruit Ripening. Plant Cell 23: 923 LP – 941. http://www.plantcell.org/content/23/3/923.abstract.

Kerckhoffs LHJ, Kelmenson PM, Schreuder MEL, Kendrick CI, Kendrick RE, Hanhart CJ, Koornneef M, Pratt LH, Cordonnier-Pratt MM. 1999. Characterization of the gene encoding the apoprotein of phytochrome B2 in tomato, and identification of molecular lesions in two mutant alleles. Mol Gen Genet 261: 901–907.

Kerckhoffs LHJ, Van Tuinen A, Hauser BA, Cordonnier-Pratt MM, Nagatani A, Koornneef M, Pratt LH, Kendrick RE. 1996. Molecular analysis of tri-mutant alleles in tomato indicates the Tri locus is the gene encoding the apoprotein of phytochrome B1. Planta 199: 152–157.

Krueger F, Andrews SR. 2011. Bismark: A flexible aligner and methylation caller for Bisulfite-Seq applications. Bioinformatics 27: 1571–1572.

Lang Z, Wang Y, Tang K, Tang D, Datsenka T, Cheng J, Zhang Y, Handa AK, Zhu J-K,. 2017. Critical roles of DNA demethylation in the activation of ripening-induced genes and inhibition of ripening-repressed genes in tomato fruit. Proc Natl Acad Sci 114: E4511–E4519.

Lazarova GI, Kerckhoffs LHJ, Brandstädter J, Matsui M, Kendrick RE, Cordonnier-Pratt MM, Pratt LH. 1998. Molecular analysis of PHYA in wild-type and phytochrome A-deficient mutants of tomato. Plant J 14: 653–662.

Lei M, Zhang H, Julian R, Tang K, Xie S, Zhu J-K,. 2015. Regulatory link between DNA methylation and active demethylation in *Arabidopsis* Proc Natl Acad Sci 112: 3553 LP – 3557. http://www.pnas.org/content/112/11/3553.abstract.

Li Z, Jiang G, Liu X, Ding X, Zhang D, Wang X, Zhou Y, Yan H, Li T, Wu K, et al. 2020. Histone demethylase SlJMJ6 promotes fruit ripening by removing H3K27 methylation of ripening-related genes in tomato. New Phytol 227: 1138–1156. https://doi.org/10.1111/nph.16590.

Liu J, Tang X, Gao L, Gao Y, Li Y, Huang S, Sun X, Miao M, Zeng H, Tian X, et al. 2012. A Role of Tomato UV-Damaged DNA Binding Protein 1 (DDB1) in Organ Size Control via an Epigenetic Manner. PLoS One 7: e42621. https://doi.org/10.1371/journal.pone.0042621.

Liu R, How-Kit A, Stammitti L, Teyssier E, Rolin D, Mortain-Bertrand A, Halle S, Liu M, Kong J, Wu C, et al. 2015. A DEMETER-like DNA demethylase governs tomato fruit ripening. Proc Natl Acad Sci 112: 10804 LP – 10809. http://www.pnas.org/content/112/34/10804.abstract.

Lu X, Wang W, Ren W, Chai Z, Guo W, Chen R, Wang L, Zhao J, Lang Z, Fan Y, et al. 2015. Genome-Wide Epigenetic Regulation of Gene Transcription in Maize Seeds. PLoS One 10: e0139582. https://doi.org/10.1371/journal.pone.0139582.

Manning K, Tör M, Poole M, Hong Y, Thompson AJ, King GJ, Giovannoni JJ, Seymour GB. 2006. A naturally occurring epigenetic mutation in a gene encoding an SBP-box transcription factor inhibits tomato fruit ripening. Nat Genet 38: 948–952. https://doi.org/10.1038/ng1841.

Mazzella MA, Arana MV, Staneloni RJ, Perelman S, Rodríguez-Batiller MJ, Muschietti J, Cerdán PD, Chen K, Sánchez RA, Zhu T, et al. 2005. Phytochrome control of the Arabidopsis transcriptome anticipates seedling exposure to light. Plant Cell 17: 2507–2516.

McCarthy DJ, Chen Y, Smyth GK. 2012. Differential expression analysis of multifactor RNA-Seq experiments with respect to biological variation. Nucleic Acids Res 40: 4288–4297. https://doi.org/10.1093/nar/gks042.

Mizrahi Y, Dostal HC, Cherry JH. 1976. Protein differences between fruits of rin, a non-ripening tomato mutant, and a normal variety. Planta 130: 223–224. https://doi.org/10.1007/BF00384424.

Omidvar V, Fellner M. 2015. DNA Methylation and Transcriptomic Changes in Response to Different Lights and Stresses in 7B-1 Male-Sterile Tomato. PLoS One 10: e0121864. https://doi.org/10.1371/journal.pone.0121864.

Paik I, Huq E. 2019. Plant photoreceptors: Multi-functional sensory proteins and their signaling networks. Semin Cell Dev Biol 92: 114–121. https://doi.org/10.1016/j.semcdb.2019.03.007.

Perrella G, Kaiserli E. 2016. Light behind the curtain: photoregulation of nuclear architecture and chromatin dynamics in plants. New Phytol 212: 908–919.

Pikaard CS, Scheid OM. 2014. Epigenetic regulation in plants. Cold Spring Harb Perspect Biol 6: 1–32.

Pontvianne F, Blevins T, Pikaard CS. 2010. Arabidopsis Histone Lysine Methyltransferases. 53: 1–22.

Robinson MD, McCarthy DJ, Smyth GK. 2009. edgeR: a Bioconductor package for differential expression analysis of digital gene expression data. Bioinformatics 26: 139–140. https://doi.org/10.1093/bioinformatics/btp616.

Shikata H, Hanada K, Ushijima T, Nakashima M, Suzuki Y, Matsushita T. 2014. Phytochrome controls alternative splicing to mediate light responses in Arabidopsis. Proc Natl Acad Sci U S A 111: 18781–18786.

Tessadori F, van Zanten M, Pavlova P, Clifton R, Pontvianne F, Snoek LB, Millenaar FF, Schulkes RK, van Driel R, Voesenek LACJ, et al. 2009. Phytochrome B and histone deacetylase 6 control light-induced chromatin compaction in Arabidopsis thaliana. PLoS Genet 5: e1000638.

Thimm O, Bläsing O, Gibon Y, Nagel A, Meyer S, Krüger P, Selbig J, Müller LA, Rhee SY, Stitt M. 2004. mapman: a user-driven tool to display genomics data sets onto diagrams of metabolic pathways and other biological processes. Plant J 37: 914–939. http://dx.doi.org/10.1111/j.1365-313X.2004.02016.x.

Ushijima T, Hanada K, Gotoh E, Yamori W, Kodama Y, Tanaka H, Kusano M, Fukushima A, Tokizawa M, Yamamoto YY, et al. 2017. Light Controls Protein Localization through Phytochrome-Mediated Alternative Promoter Selection. Cell 171: 1316–1325.e12.

Vrebalov J, Ruezinsky D, Padmanabhan V, White R, Medrano D, Drake R, Schuch W, Giovannoni J. 2002. A MADS-Box Gene Necessary for Fruit Ripening at the Tomato *Ripening-Inhibitor*(*Rin*) Locus. Science (80−) 296: 343 LP – 346. http://science.sciencemag.org/content/296/5566/343.abstract.

Yang C, Shen W, Yang L, Sun Y, Li X, Lai M, Wei J, Wang C, Xu Y, Li F, et al. 2020. HY5-HDA9 Module Transcriptionally Regulates Plant Autophagy in Response to Light-to-Dark Conversion and Nitrogen Starvation. Mol Plant 13: 515–531. https://doi.org/10.1016/j.molp.2020.02.011.

Zemach A, Kim MY, Hsieh PH, Coleman-Derr D, Eshed-Williams L, Thao K, Harmer SL, Zilberman D. 2013. The arabidopsis nucleosome remodeler DDM1 allows DNA methyltransferases to access H1-containing heterochromatin. Cell 153: 193–205. http://dx.doi.org/10.1016/j.cell.2013.02.033.

Zhang B, Tieman DM, Jiao C, Xu Y, Chen K, Fei Z, Giovannoni JJ, Klee HJ. 2016. Chilling-induced tomato flavor loss is associated with altered volatile synthesis and transient changes in DNA methylation. Proc Natl Acad Sci 113: 12580 LP – 12585. http://www.pnas.org/content/113/44/12580.abstract.

Zhang H, Lang Z, Zhu JK. 2018. Dynamics and function of DNA methylation in plants. Nat Rev Mol Cell Biol 19: 489–506. http://dx.doi.org/10.1038/s41580-018-0016-z.

Zhang Y, Liu T, Meyer CA, Eeckhoute J, Johnson DS, Bernstein BE, Nusbaum C, Myers RM, Brown M, Li W, et al. 2008. Model-based Analysis of ChIP-Seq (MACS). Genome Biol 9: R137. https://doi.org/10.1186/gb-2008-99-r137.

Zhao L, Lu J, Zhang J, Wu P-Y, Yang S, Wu K. 2015. Identification and characterization of histone deacetylases in tomato (Solanum lycopersicum). Front Plant Sci 5: 760. https://www.frontiersin.org/article/10.3389/fpls.2014.00760.

Zhong S, Fei Z, Chen YR, Zheng Y, Huang M, Vrebalov J, McQuinn R, Gapper N, Liu B, Xiang J, et al. 2013. Single-base resolution methylomes of tomato fruit development reveal epigenome modifications associated with ripening. Nat Biotechnol 31: 154–159. http://dx.doi.org/10.1038/nbt.2462.

Zuo J, Grierson D, Courtney LT, Wang Y, Gao L, Zhao X, Zhu B, Luo Y, Wang Q, Giovannoni JJ. 2020. Relationships between genome methylation, levels of non-coding RNAs, mRNAs and metabolites in ripening tomato fruit. Plant J.

