## Supplementary figures and images for "THE REGULATORY EFFECT OF LIGHT OVER FRUIT DEVELOPMENT AND RIPENING IS MEDIATED BY EPIGENETIC MECHANISMS"

### Supplemental Figure 1

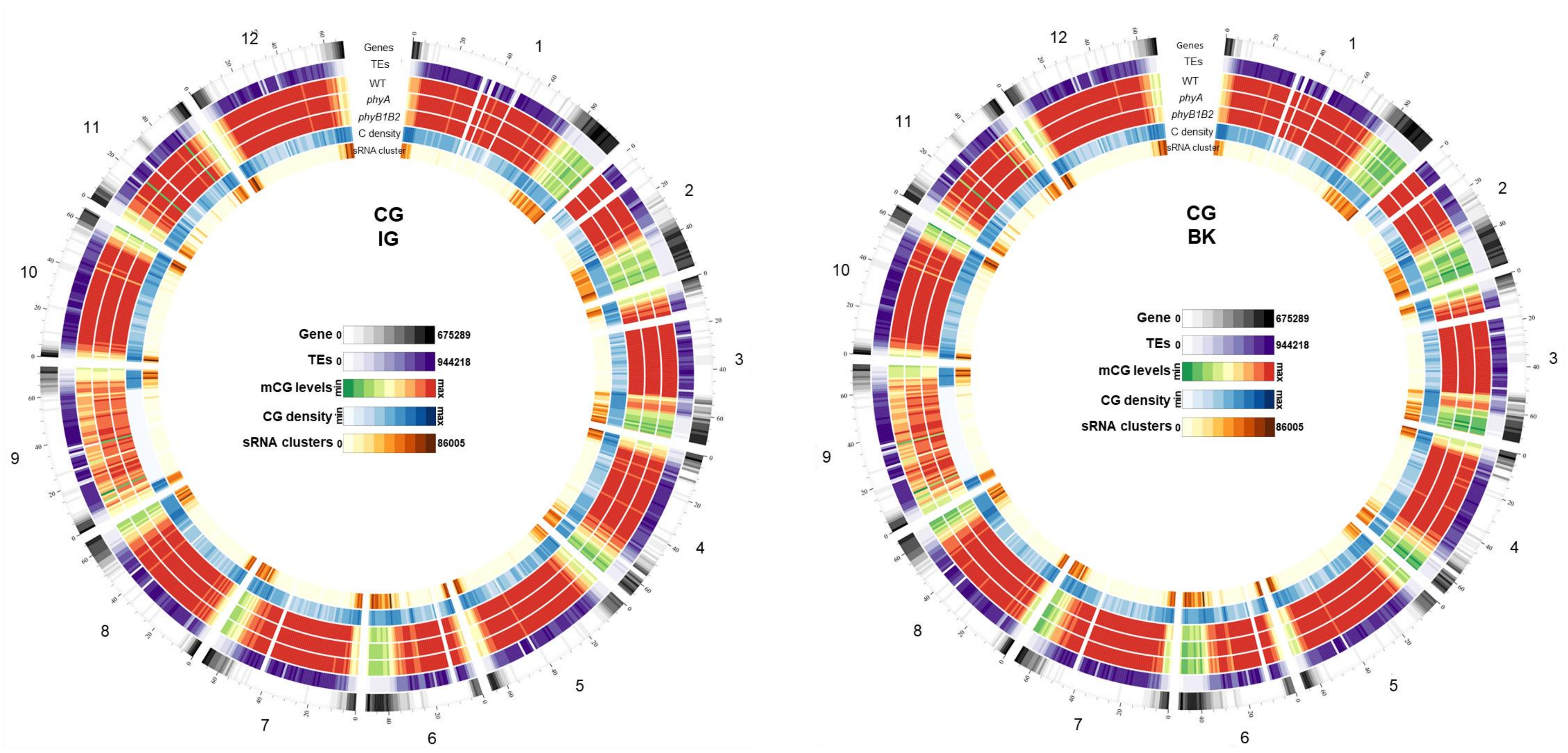

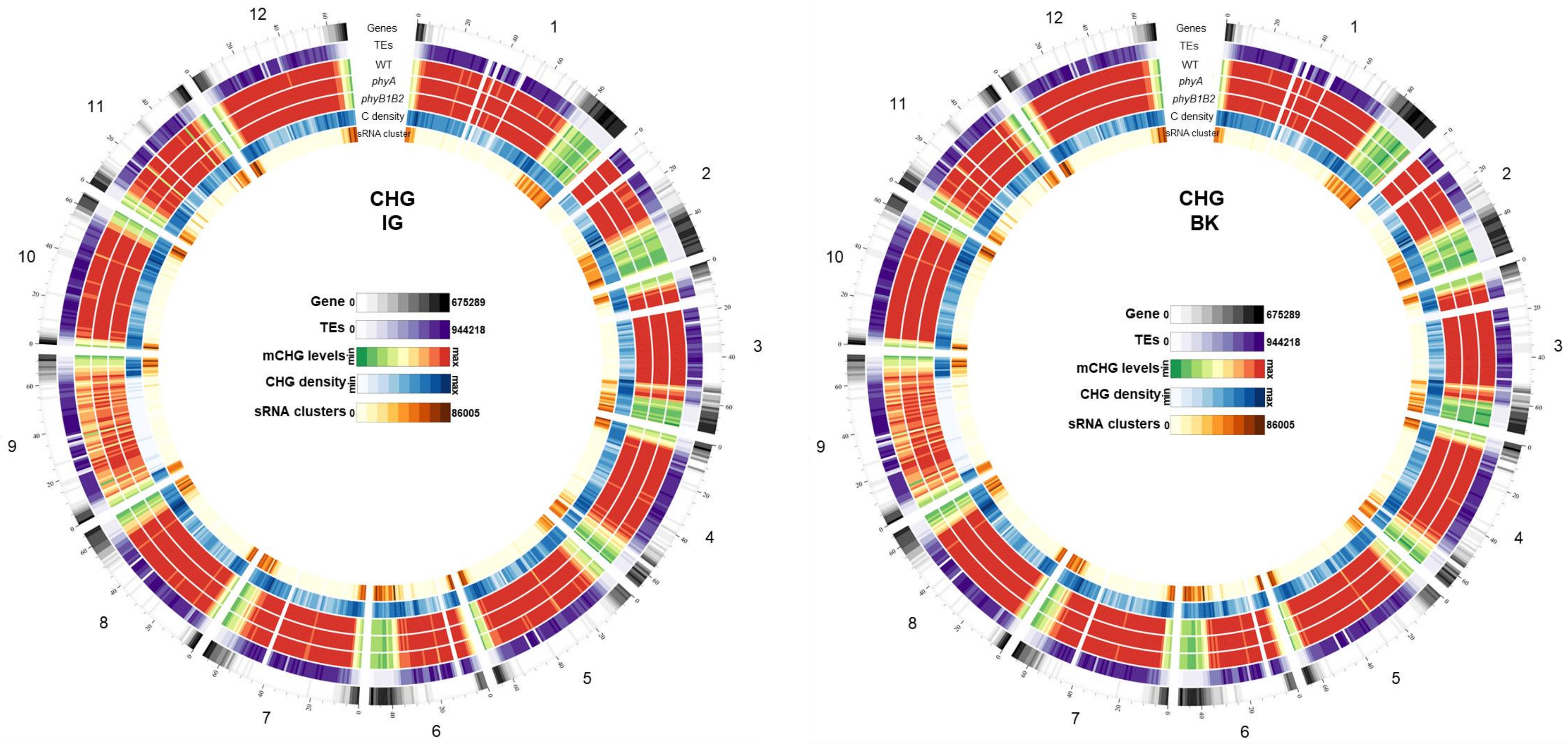

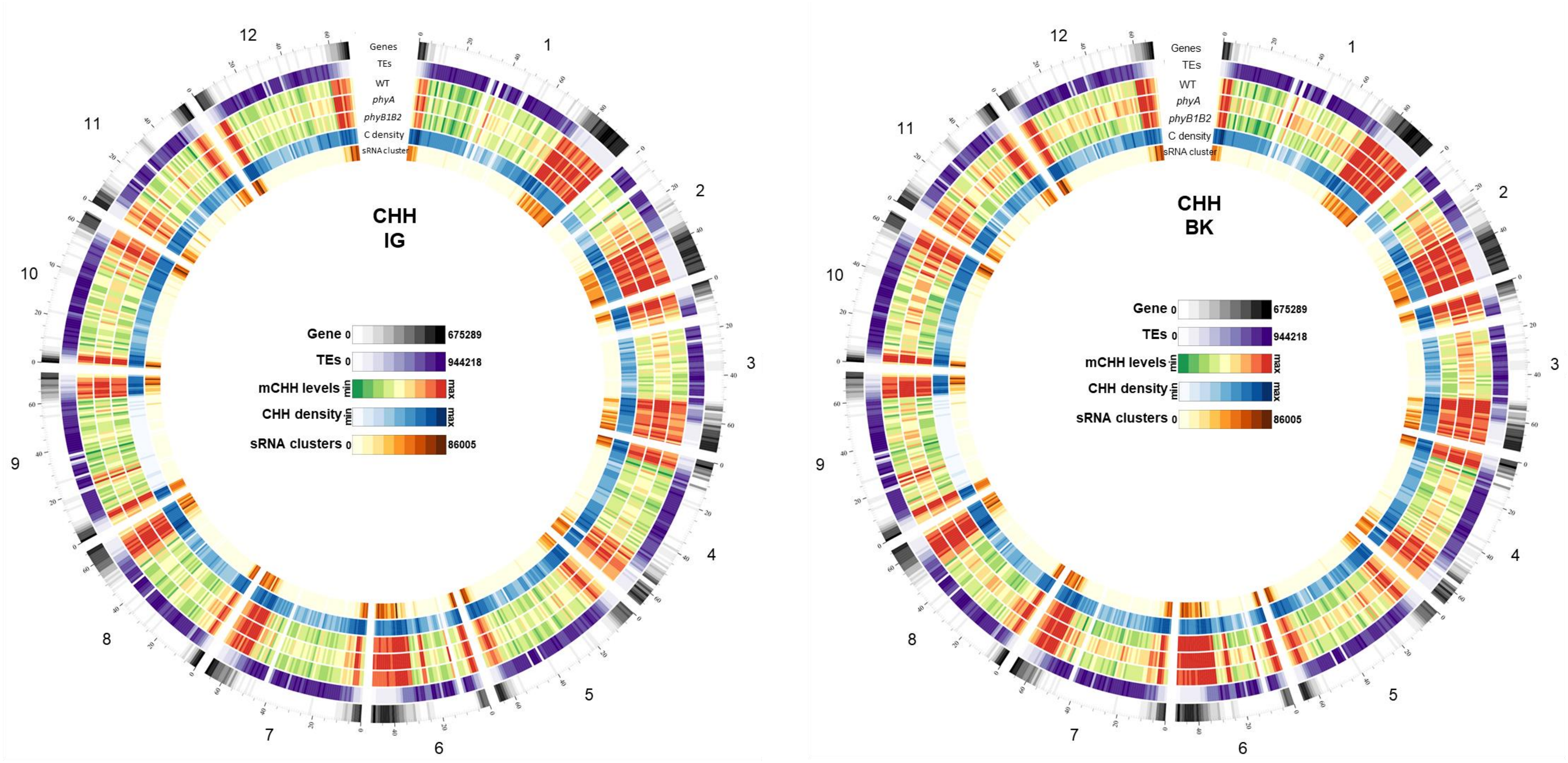

### Supplemental Figure 4

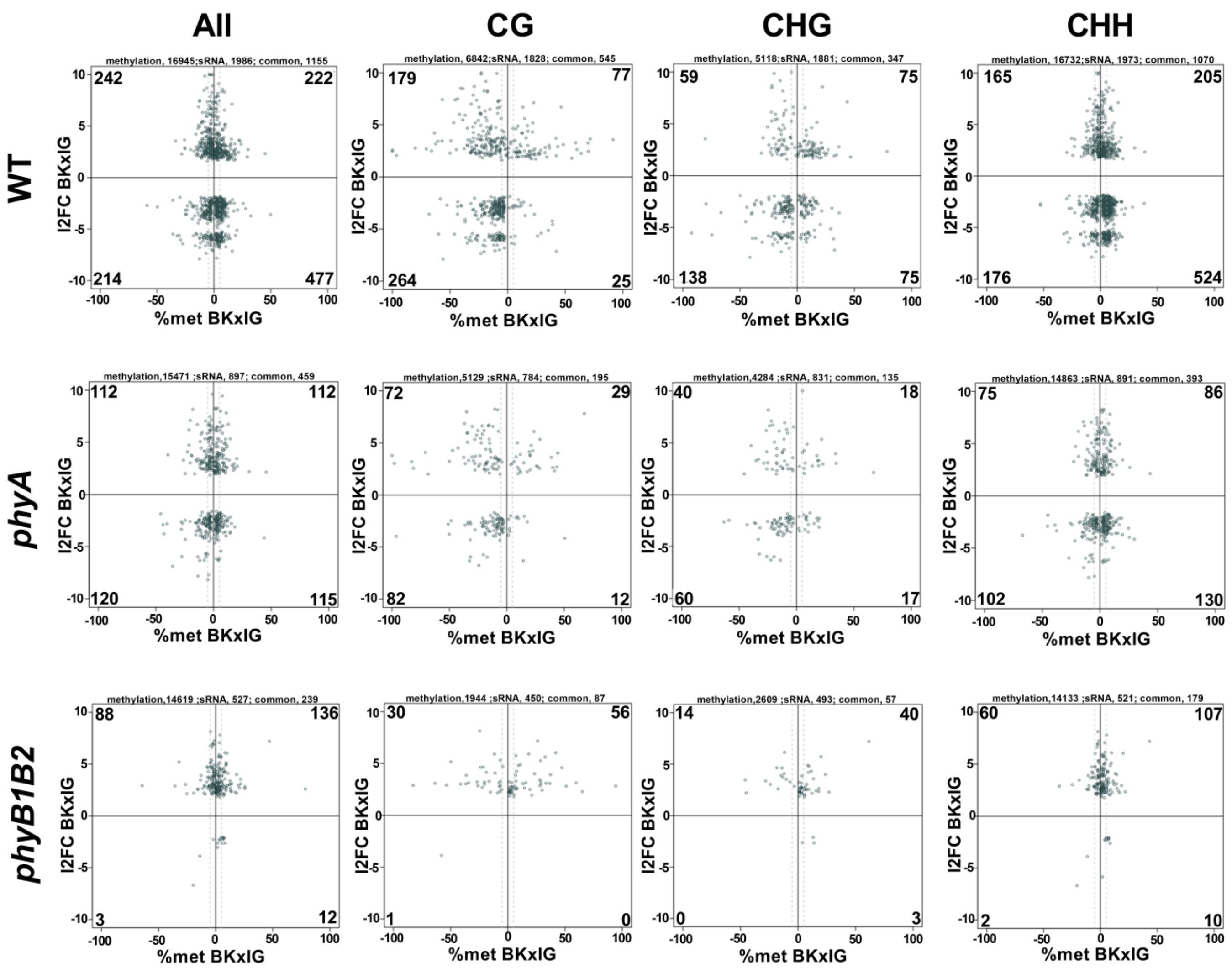

### Supplemental Figure 5

a

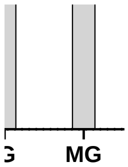


b

### Supplemental Figure 6

**a**

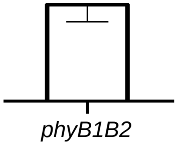


**b**

**c**

### Supplementary Figure 2

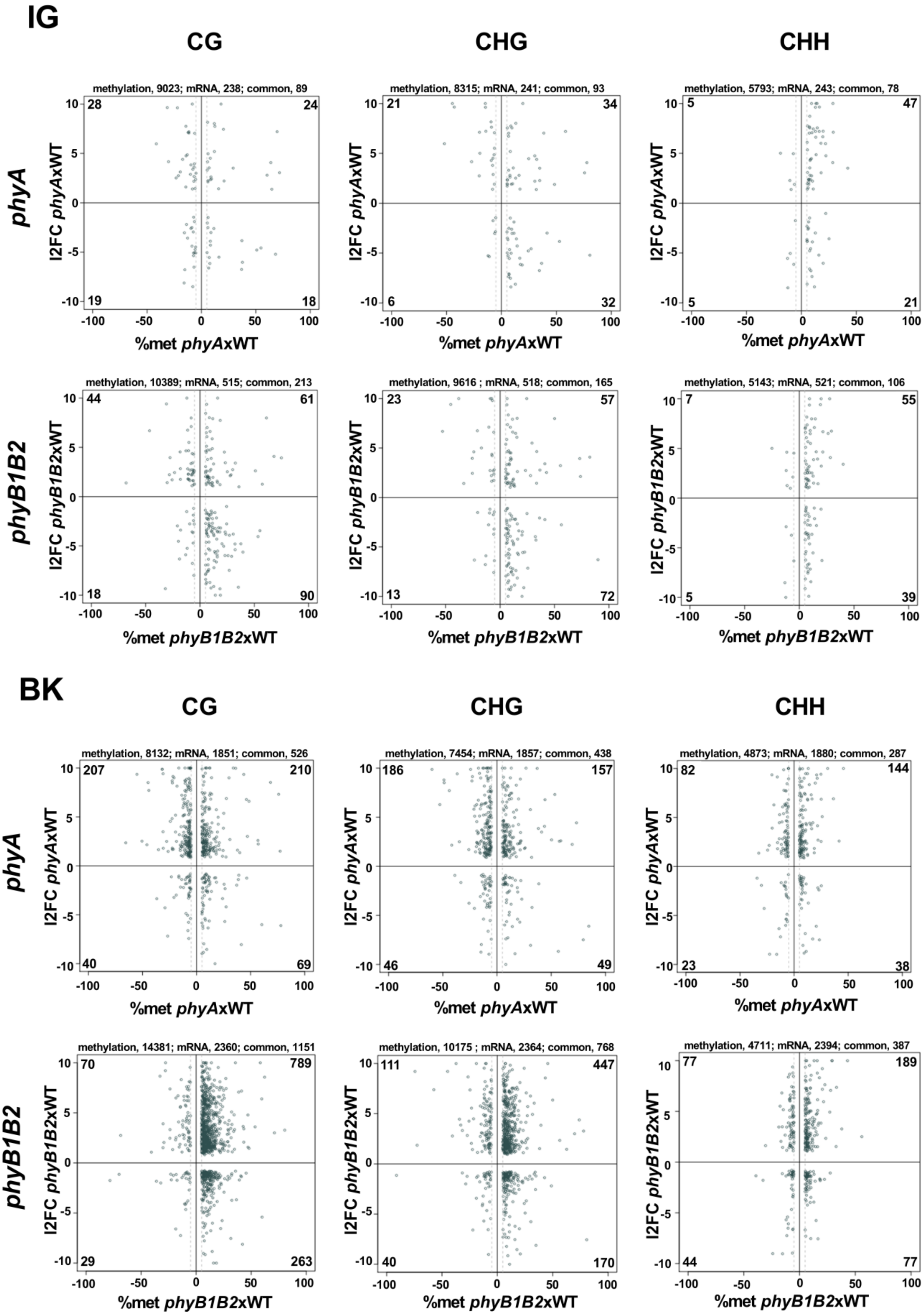

### Supplementary Figure 3

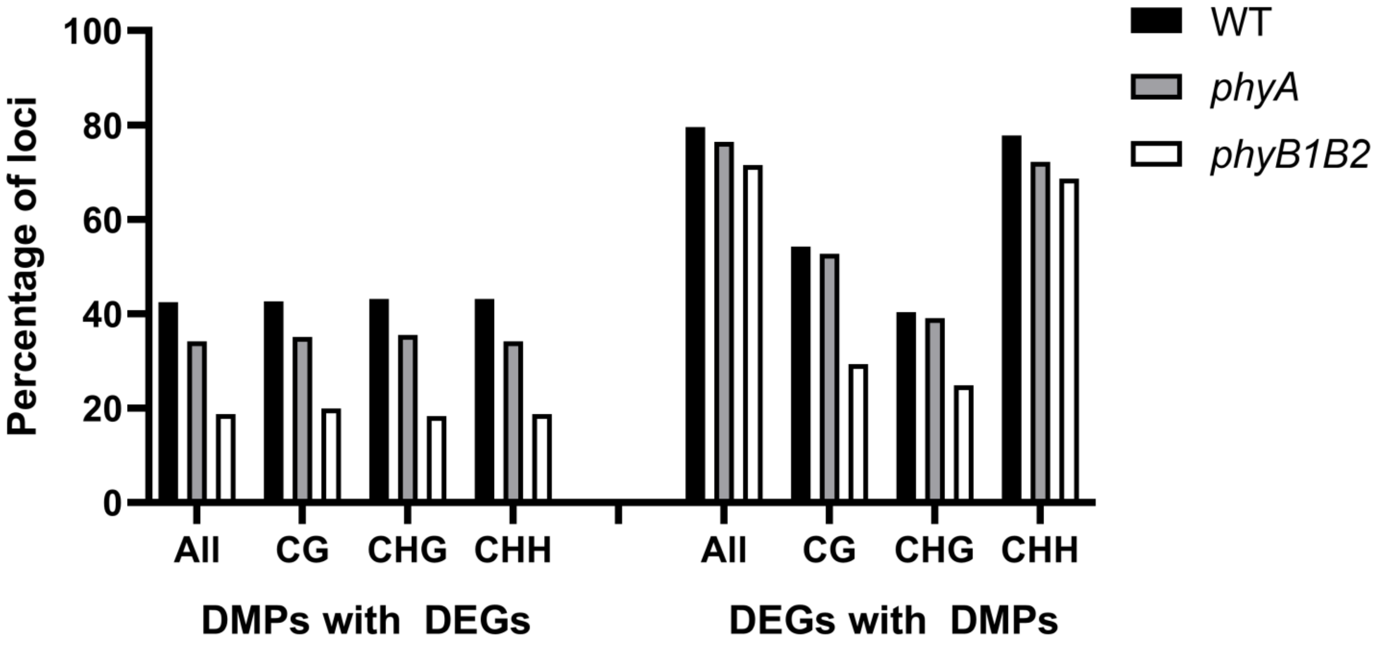

### Supplementary Figure 7

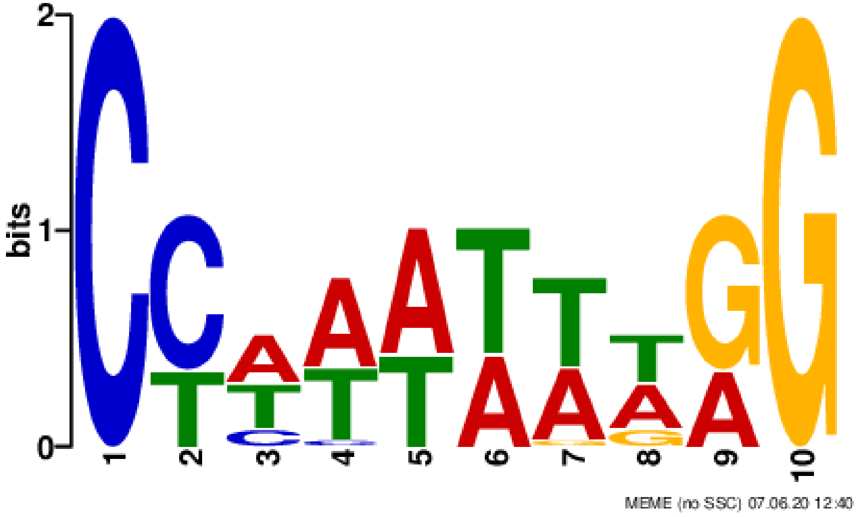

### Supplementary Figure 8

**Solyc09g00980 *ROS1L***

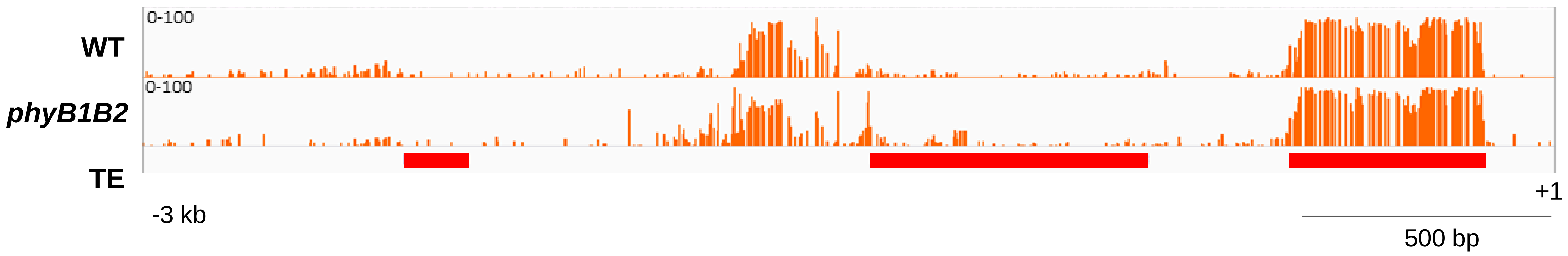
